# Sleep spindles coordinate corticostriatal reactivations during the emergence of automaticity

**DOI:** 10.1101/2020.10.25.354282

**Authors:** S. M. Lemke, D. S. Ramanathan, D. Darevsky, D. Egert, J. D. Berke, K. Ganguly

## Abstract

Plasticity within the corticostriatal network is known to regulate the balance between behavioral flexibility and automaticity. Repeated training of an action has been shown to bias behavior towards automaticity, suggesting that training may trigger activity-dependent corticostriatal plasticity. However, surprisingly little is known about the natural activity patterns that may drive plasticity or when they occur during long-term training. Here we chronically monitored neural activity from primary motor cortex (M1) and the dorsolateral striatum (DLS) during both training and offline periods, i.e., time away from training including sleep, throughout the development of an automatic reaching action. We first show that blocking striatal NMDA receptors during offline periods prevents the emergence of behavioral consistency, a hallmark of automaticity. We then show that, throughout the development of an automatic reaching action, corticostriatal functional connectivity increases during offline periods. Such increases track the emergence of consistent behavior and predictable cross-area neural dynamics. We then identify sleep spindles during non-REM sleep (NREM) as uniquely poised to mediate corticostriatal plasticity during offline periods. We show that sleep spindles are periods of maximal corticostriatal transmission within offline periods, that sleep spindles in post-training NREM reactivate neurons across areas, and that sleep-spindle modulation in post-training NREM is linked to observable changes in spiking relationships between individual pairs of M1 and DLS neurons. Our results indicate that offline periods, in general, and sleep spindles, specifically, play an important role in regulating behavioral flexibility through corticostriatal network plasticity.

## Introduction

Automaticity allows animals to capitalize on invariance in the environment through the development of actions that, while inflexible to changes, are performed highly consistently in response to a specific stimulus^1–5^. It has been demonstrated that the consistent production of an action emerges with repeated training over multiple days and is accompanied by coordinated neural activity across the corticostriatal network during action execution^6–10^. Importantly, the emergence of such actions has also been shown to require striatal NMDA receptor activation^10–12^, suggesting that cortical activity pattens that modulate the striatum may be important drivers of activity-dependent plasticity^13,14^ and the emergence of coordinated corticostriatal activity. However, surprisingly little is known about the natural activity patterns related to repeated task training that underlie the emergence of automaticity.

One intriguing possibility is that time away from training - “offline” periods, including sleep - may play a role in modifying the corticostriatal network. This possibility is motivated by evidence that sleep-dependent reactivations of cortical neural ensembles active during task performance are essential for initial learning^15–18^. It is possible that the coordinated reactivations of both cortical and striatal ensembles modify the corticostriatal network and impact behavior during long-term training^19–22^. However, how cortical reactivation events engage downstream striatal ensembles remains unclear. Moreover, how such cross-area activity may precisely modify the corticostriatal network and impact network activity during subsequent awake behavior is unexplored.

Currently, our understanding of how sleep impacts distributed brain networks is largely derived from the *systems consolidation theory*, where it has been shown that coordinated activity patterns across hippocampus and cortex lead to the formation of stable long-term memories in cortex that do not require the hippocampus^23–25^. Notably, whether sleep impacts the connectivity across hippocampus and cortex has not been established. Therefore, one possibility is that, in the corticostriatal network, we similarly observe coordinated cross-area activity patterns during sleep but do not find evidence for the modification of corticostriatal connectivity during offline periods. Alternatively, it is possible that we find evidence that cross-area activity patterns during sleep modify the connectivity between cortex and striatum and impact network activity during subsequent behavior.

Here we establish that offline periods play an essential role in modifying the corticostriatal network during the emergence of automaticity and identify sleep spindles as uniquely poised to mediate such plasticity. We show that during post-training NREM, sleep-dependent reactivation events are coordinated across both cortex and striatum during sleep spindles and link such spindle-modulation to changes in functional connectivity across the corticostriatal network. These results suggest that sleep plays an important role in modifying cross-area connectivity within offline periods and that the modulation of activity patterns during sleep may offer novel therapeutic targets for unlearning maladaptive habits^26,27^.

## Results

To study how corticostriatal network activity evolves during long-term training, we implanted six adult rats with either microwire electrode arrays (*n* = 4) or custom built high-density silicon probes^28^ (*n* = 2) in both primary motor cortex (M1) and the dorsolateral striatum (DLS), which receives the majority of M1 projections to the striatum^29^ (**Figure 1a**). Neural activity across regions was monitored as rats underwent ~eight days of reach-to-grasp task training (range: 5-14 days, mean: 8.67 days). Each recording day consisted of a 2-3 hour pre-training block (“pre-sleep”), a 100-150 trial training block, and a second 2-3 hour post-training block (“post-sleep”; **Figure 1b**; pre-sleep length: 157.2 ± 5.8 minutes, post-sleep length: 166.6 ± 6.3 minutes, mean ± SEM). The reach-to-grasp task requires rats to reach and grasp a food pellet through a small window present in their behavioral box. During pre- and post-sleep, behavioral states, i.e., wake, non-REM sleep (NREM), and REM sleep, were classified using standard methods based on cortical local field potential (LFP) power and movement measured from video or electromyography (EMG) activity^30^.

**Figure 1.**
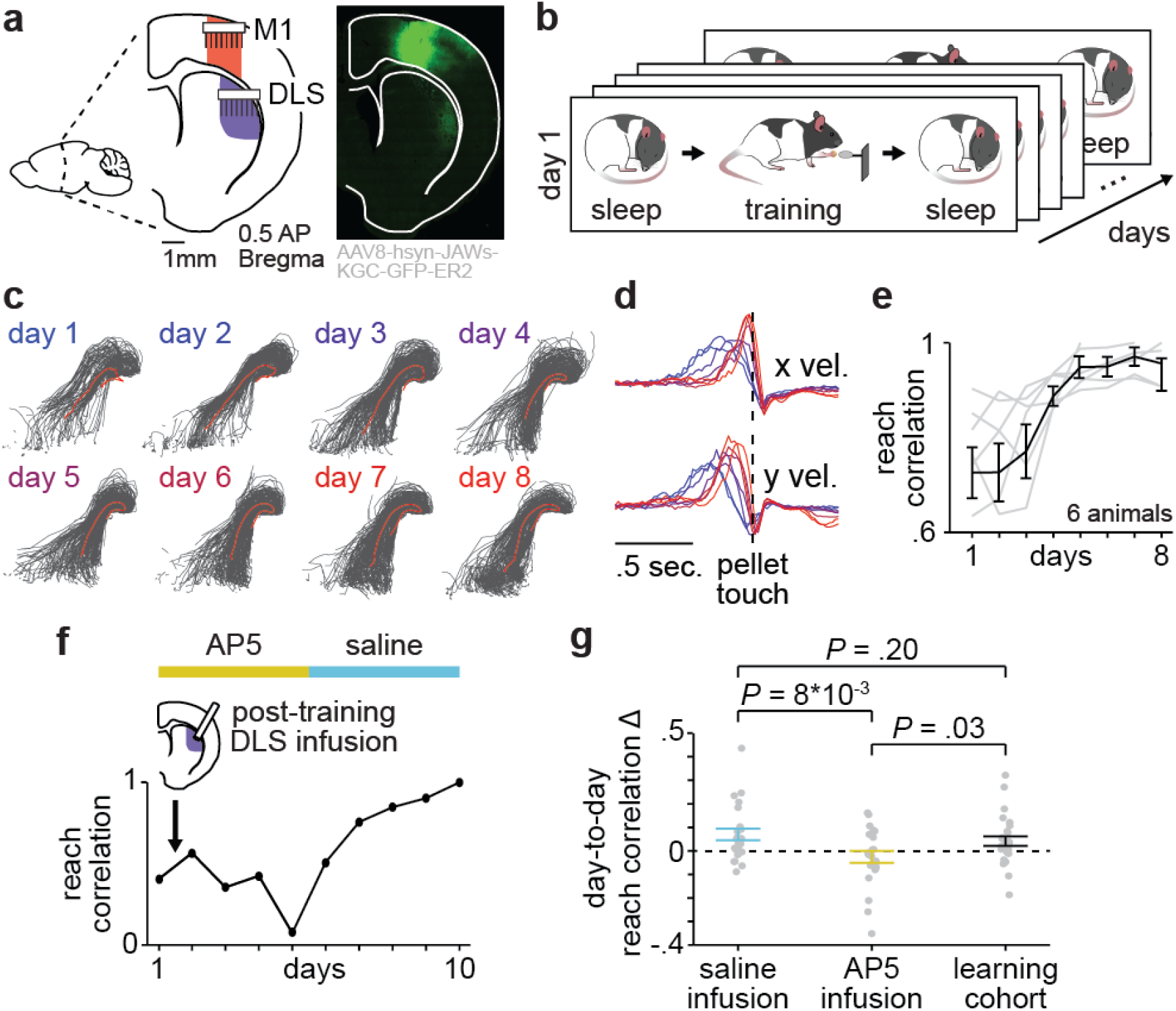
Offline striatal NMDA receptor activation is required to develop a consistent behavior. **a.** Schematic displaying primary motor cortex (M1) and dorsolateral striatum (DLS) recording locations (left) and labeled M1 projections showing direct input to the DLS (right). **b.** Schematic showing each day’s recording blocks during long-term training. **c.** Individual reach trajectories in grey overlaid with mean reach trajectory across trials in red for each day of training in example animal. **d.** Average reach velocity profile in x and y dimensions for each day of training in example animal. **e.** Reach velocity profile correlation for first eight days of training for individual animals in grey overlaid with mean ± SEM across animals in black. **f.** Day-to-day evolution in reach velocity profile correlation with post-training DLS infusions of either AP5 or saline in example animal. **g.** Comparison of day-to-day changes in reach velocity profile correlation with post-training saline infusion, post-training AP5 infusion, or no infusion in learning cohort animals, showing specific decrease in day-to-day reach velocity profile correlation with post-training AP5 infusion. Individual day-to-day changes as grey dots overlaid with mean ± SEM across all day-to-day changes in color.

### Offline striatal NMDA receptor activation is required to develop a consistent behavior

With repeated training on the reach-to-grasp task, animals developed a consistent reaching trajectory (**Figure 1c**) and reaching velocity profile (**Figure 1d**). Measuring the correlation between the mean reaching velocity profile on each day of training and the final day of training revealed that a consistent day-to-day reaching action emerged within the first eight days of training (**Figure 1e**). Such day-to-day invariance in skilled reaching is consistent with the emergence of automaticity^1–3,31^. To further test the automaticity of reaching after the emergence of invariant behavior, we moved the location of the food pellet such that reaches to the old pellet location would no longer be successful. If an animal were reaching flexibly, we would expect that reaching behavior would quickly adapt to the new position. Alternatively, if an animal were reaching automatically, we would expect that reaches would remain consistent despite decreased success. Consistent with automaticity, reach trajectories remained consistent and did not adapt to the new pellet position despite a large decrease in success rate (**Supplemental Figure 1**). We also examined whether such automaticity emerged with long-term training or existed at the start of training by testing whether animals could reach flexibility during the first two days of exposure to the task (*n* = 2 rats with no neural implant). These animals were able to reach to each of the two different pellet positions with comparable success rates (animal 1: 43% and 72% success rate; animals 2: 44% and 50% success rate, ~200 trials in each animal), indicating that their reaching behavior was flexible prior to repeated training. Altogether this indicated that our training paradigm led to the formation of inflexible and automatic reach-to-grasp behavior.

To further test the link between offline plasticity in the corticostriatal network and increases in behavioral consistency, we trained a new cohort of animals (*n* = 6 rats) and infused 1μl of either NMDA receptor antagonist AP5 (5μg/μl) or saline into DLS immediately after training on each day (**Figure 1f**). This revealed that offline striatal NMDA activation was essential for the emergence of a consistent reaching behavior, as day-to-day changes in reach consistency were significantly decreased with AP5 infusions, compared to saline infusions or day-to-day improvements observed in the learning cohort (**Figure 1g**; *n* = 24 correlation change values with AP5 infusions, −0.03 ± 0.03 correlation value, *n* = 24 correlation change values with saline infusions, 0.07 ± 0.02 correlation value, *n* = 40 correlation change values in learning cohort, 0.04 ± 0.01 correlation value; AP5 infusions vs. saline infusions: *t*(23) = 2.8, *P* = 8×10^-3^, paired-sample t-test, AP5 infusions vs. learning cohort: *t*(62) = 2.3, *P* = 0.03, two-sample t-test, saline infusions vs. learning cohort: *t*(62) = 1.3, *P* = 0.20, two-sample t-test). Importantly, task engagement, as measured by reaction time from trial start to reach, did not differ for trials on subsequent days after AP5 or saline infusions (AP5: 218.2 ± 3.2ms, saline: 222.4 ± 3.0ms, *t*(5198) = −0.96, *P* = 0.34, two-sample t-test). Altogether, these results were consistent with the notion that offline plasticity in the corticostriatal network following training is critical for the emergence of automaticity.

### Corticostriatal functional connectivity increases during offline periods

To measure long-term changes in corticostriatal functional connectivity during the emergence of automaticity, we measured LFP coherence across individual pairs of M1 and DLS electrodes. LFP signals can be stably recorded across multiple days allowing LFP coherence to provide a stable long-term measure of multi-region connectivity^32,33^. Specifically, within the corticostriatal network, theta coherence (4-8Hz) has been previously shown to reflect coordinated population spiking activity^8,9,34^. Therefore, we measured 4-8Hz LFP coherence during pre- and post-sleep on each day of training to determine when corticostriatal functional connectivity changed during long-term training (**Figure 2a**). LFP coherence was calculated specifically during NREM to control for any differences in the time spent in each behavioral state during pre- and post-sleep. Common-mode referencing was applied, separately in each region, to decrease common noise and minimize volume conduction^8^. We found that there was a significant correlation between each day’s mean 4-8Hz LFP coherence across all channel pairs and reach velocity profile correlation (r = 0.44, *P* = 7×10^-3^, Pearson’s *r*), indicating that offline LFP coherence reflects changes in corticostriatal functional connectivity that are related to the emergence of a consistent behavior.

**Figure 2.**
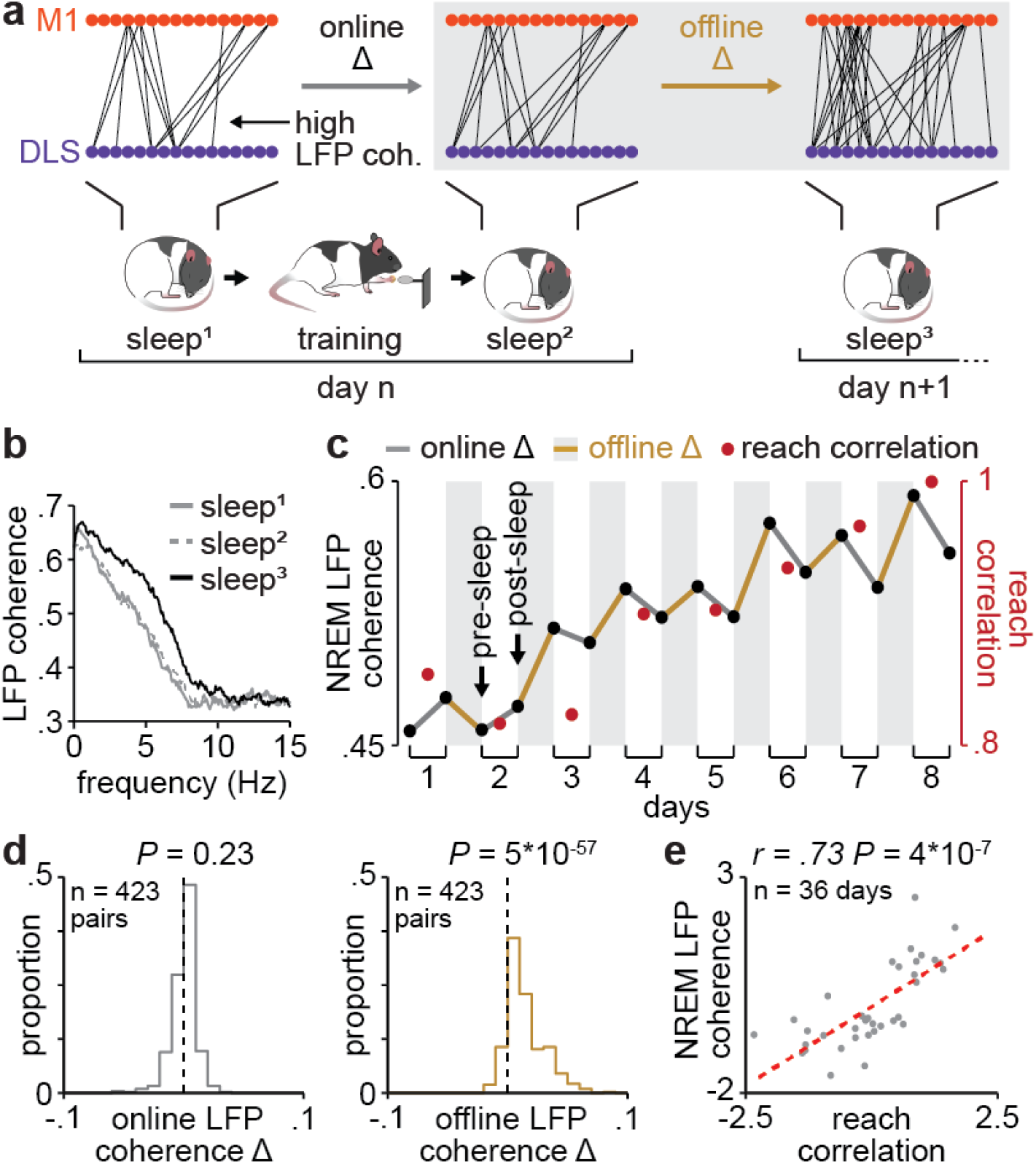
Corticostriatal functional connectivity increases during offline periods. **a.** Schematic depicting M1 and DLS electrode pairs with high 4-8Hz LFP coherence (>0.6 coherence value measured in NREM) during pre- and post-sleep on one day of training and pre-sleep on the next day of training, showing an increase in the number of high LFP coherence pairs occurring offline rather than online, in example animal. **b.** LFP coherence spectrums (measured in NREM) across example M1 and DLS electrode pair for pre- and post-sleep periods represented in panel **a.** showing an increase in 4-8Hz LFP coherence largely occurring offline rather than online. **c.** LFP coherence (4-8Hz measured in NREM) for each pre- and post-sleep period throughout learning for example M1 and DLS electrode pair, showing increases in coherence largely occurring offline rather than online, overlaid with reach velocity profile correlation values for each day of training. **d.** Comparison of distributions of online (left) and offline (right) changes in LFP coherence (4-8Hz measured in NREM) averaged across training days for M1 and DLS electrode pairs across animals. **e.** Correlation between each day’s mean LFP coherence (mean 4-8Hz measured in NREM during both pre- and post-sleep) and reach velocity profile correlation value.

We next sought to determine whether LFP coherence increased during training or offline periods. To do this, we specifically examined LFP channel pairs that increased in coherence from day one to day eight (33% of pairs increased, 17% decreased, and 50% did not change; increase or decrease defined as a change in coherence of at least 0.25). Remarkably, within the subset of channels that showed training-related increases in coherence over learning, increases occurred largely offline, i.e., between each day’s post-sleep and the next day’s pre-sleep, rather than online during training, i.e., between pre- and post-sleep on the same day (**Figure 2b&c**). More specifically, the distribution of online LFP coherence changes was not significantly different than zero, while the distribution of offline LFP coherence changes was skewed toward larger increases (**Figure 2d**; online LFP coherence changes: *t*(422) = 1.2, *P* = 0.23, offline LFP coherence changes: *t*(422) = 18.8, *P* = 5×10^-57^, one-sample t-test). Importantly, the subset of channels that showed training-related increases had a close relationship to the emergence of consistent behavior (**Figure 2e**; r = 0.73, *P* = 4×10^-7^, Pearson’s *r*), providing evidence that offline increases in corticostriatal functional connectivity are relevant to the consistency of behavior during the emergence of automaticity.

### Offline increases in functional connectivity predict the emergence of low-dimensional cross-area neural dynamics during behavior

We next examined how offline increases in corticostriatal functional connectivity may impact corticostriatal network activity during subsequent reach-to-grasp performance. We extracted low-dimensional neural trajectory representations of DLS spiking activity during reaching using principle components analysis (PCA). We then examined the evolution of how spiking activity in M1 could predict DLS neural trajectories over the course of training (**Figure 3a**). We found that the ability to predict DLS neural trajectories during reaching from M1 spiking activity increased with training, while the ability to predict the trajectory representations of DLS activity during a baseline, non-reaching, period did not significantly change (**Figure 3b**; reach activity: first two days of training: 0.15 ± 0.05 Pearson’s *r*, last two days of training: 0.46 ± 0.05 Pearson’s *r*, *t*(30) = −4.4, *P* = 1×10^-4^, two-sample t-test; baseline activity: first two days of training: 0.03 ± 0.02 Pearson’s *r*, last two days of training: 0.01 ± 0.03 Pearson’s *r*, *t*(30) = 1.0, *P =* 0.30, two-sample t-test). Notably, the ability to predict DLS neural trajectories during reaching from M1 spiking activity was significantly correlated to the mean 4-8Hz LFP coherence measured offline on each day of training (**Figure 3c**), indicating that offline increases in LFP coherence track the emergence of predictable cross-area dynamics during subsequent reach-to-grasp performance.

**Figure 3.**
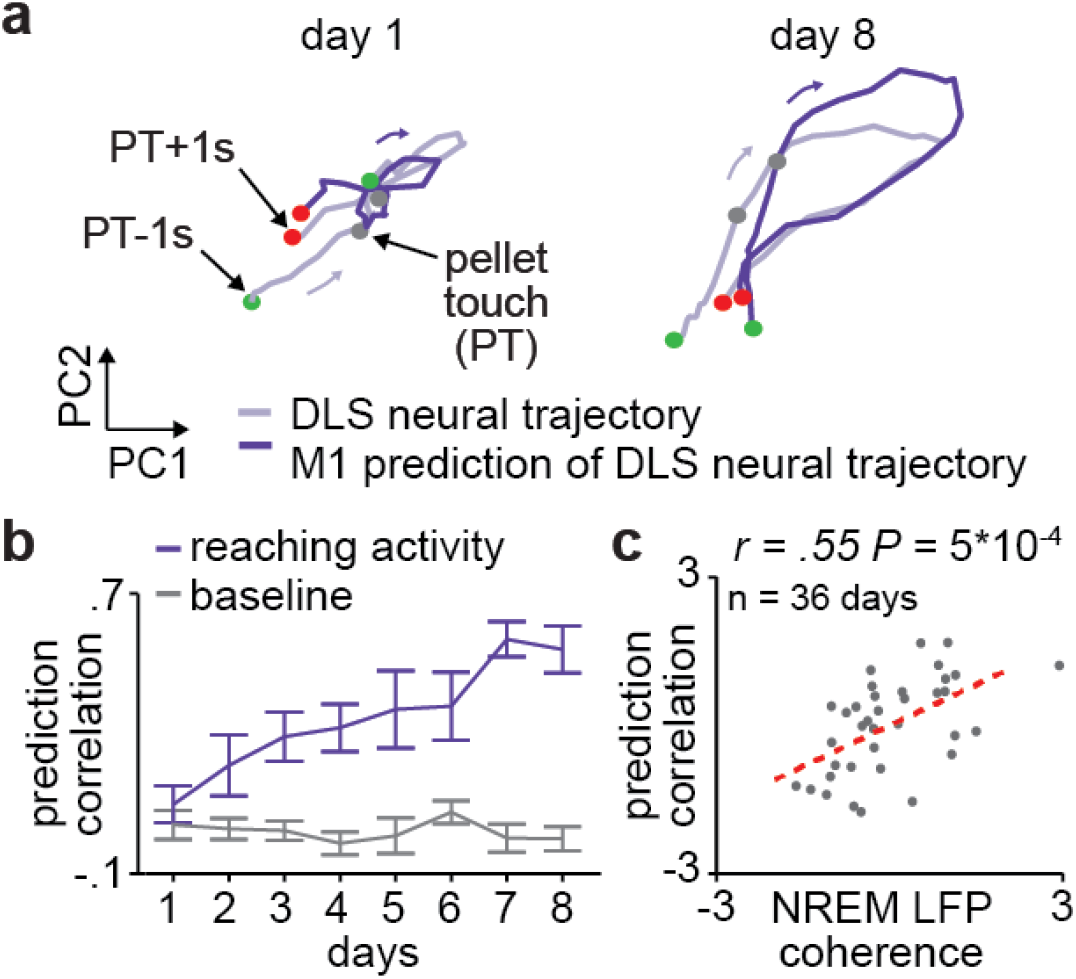
Offline increases in functional connectivity predict the emergence of low-dimensional cross-area neural dynamics during behavior. **a.** Trial-averaged neural trajectory (PC1 and PC2) of DLS activity during reaching (one second before to one second after pellet touch) on day one (left) and day eight (right) of training in example animal, overlaid with prediction of DLS neural trajectory from M1 spiking activity. **b.** Ability to predict DLS neural trajectory (PC1 and PC2) during reaching and during a baseline, non-reaching, period from M1 spiking activity on each day of training (mean ± SEM across animals). **c.** Correlation between each day’s mean LFP coherence (mean 4-8Hz measured in NREM during both pre- and post-sleep) and ability to predict DLS neural trajectory (PC1 and PC2) during reaching from M1 spiking activity.

### Corticostriatal transmission strength within offline periods is maximal during sleep spindles in NREM

Given the evidence that offline periods are relevant for changes in corticostriatal functional connectivity, we next sought to identify the activity patterns that may be responsible for driving such plasticity across M1 and DLS. To do this, we first examined how corticostriatal transmission strength, i.e., the degree to which M1 neural activity drives DLS activity, differed across behavioral states during offline periods (**Figure 4a**). To measure this, we characterized putative monosynaptically connected pairs of M1 and DLS units (*n* = 1,100 M1 and 579 DLS units) by determining whether there was a significant peak in the cross correlation of their spiking activity at the short-latency time lag consistent with the conduction and synaptic delays between M1 and DLS (~6ms time lag from M1 to DLS activity^9^; **Figure 4b**; 3,969/10,286 M1 and DLS unit pairs were classified as putatively connected; **Supplemental Figure 2a&b**). We then compared the short-latency cross correlation magnitude (1-10ms time lag) for the population of putatively connected M1 and DLS pairs across behavioral states. To account for differences in firing rates across behavioral states (**Supplemental Figure 2c&d**), we normalized each pair’s cross correlation by the mean cross correlation value from 50-100ms time lag, where no consistent spiking relationship is expected between putatively connected pairs of M1 and DLS units. This revealed that corticostriatal transmission strength was maximal during NREM, compared to REM or wake (**Figure 4c & Supplemental Figure 3a&b**).

**Figure 4.**
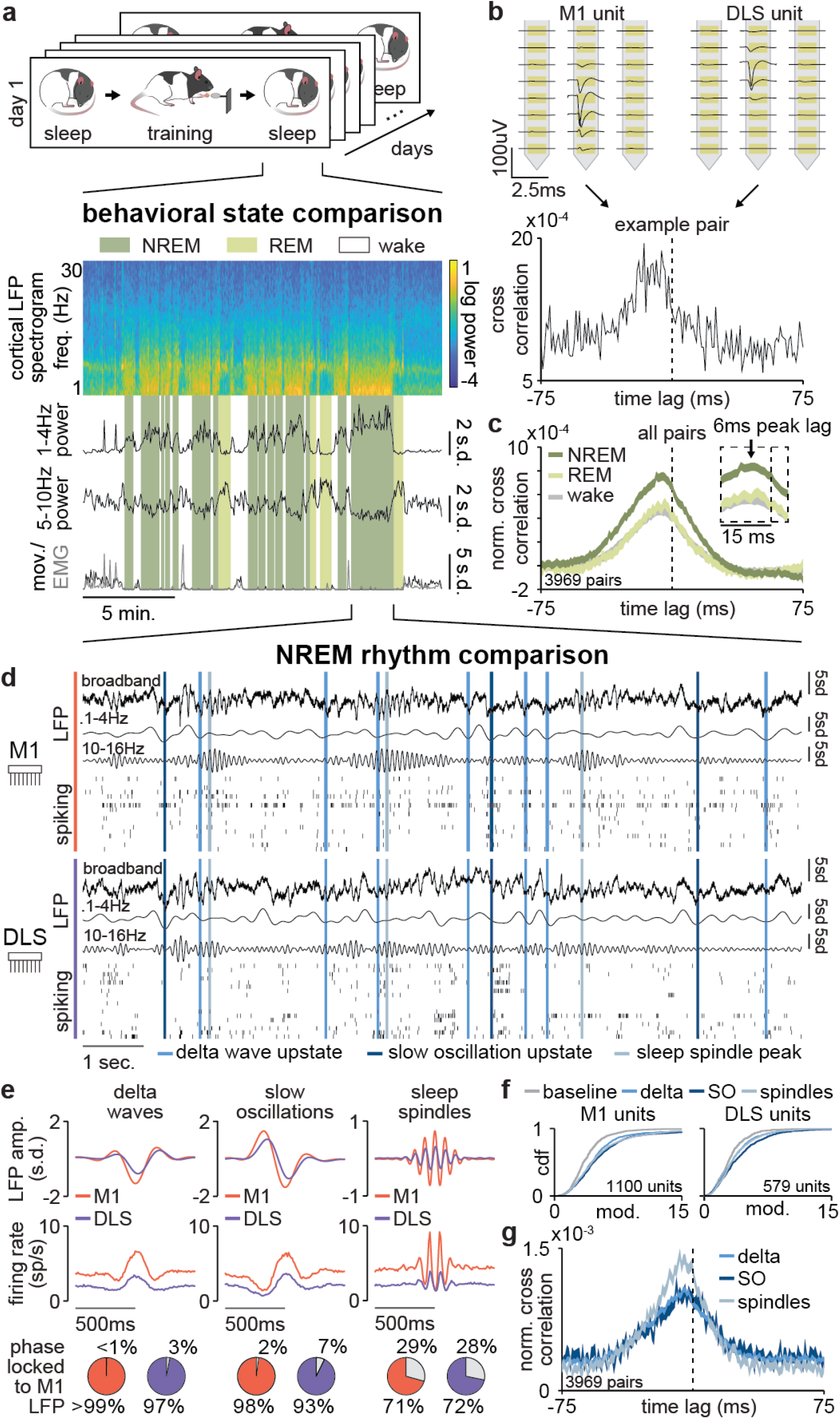
Corticostriatal transmission strength within offline periods is maximal during sleep spindles in NREM. **a.** M1 local field potential (LFP) spectrogram and behavioral state detection from example session. **b**. Example M1 and DLS single unit sorting from high-density silicon probe (top) and cross correlation of spiking activity centered on DLS unit spiking for an example pair of M1 and DLS units showing a short-latency peak indicating putative monosynaptically connectivity (bottom). **c.** Comparison of normalized cross correlations of spiking activity from all putatively connected pairs of M1 and DLS units across behavioral states, showing that corticostriatal transmission strength is maximal in NREM (width of line represents mean ± SEM). **d.** Snippet of LFP and single unit spiking activity from M1 and DLS during NREM overlaid with detected NREM rhythms in M1. **e.** Mean LFP and spiking activity during slow oscillations, delta waves, and sleep spindles in both M1 and DLS in example animal (top) and percentage of M1 and DLS units across animals significantly phase locked to M1 LFP during each NREM rhythm (significance threshold of *P* = 0.05, Rayleigh test of uniformity, bottom). **f.** Comparison of firing rate modulation distributions for M1 (left) and DLS (right) units across animals during slow oscillations, delta waves, sleep spindles, and a baseline NREM period. **g.** Comparison of normalized cross correlations of spiking activity for all putatively connected pairs of M1 and DLS units across NREM rhythms, showing that corticostriatal transmission strength is maximal during sleep spindles (width of line represents mean ± SEM).

Given the heterogeneous nature of NREM activity, we next explored the dynamics of corticostriatal transmission within NREM. We specifically detected NREM rhythms in M1 that have been previously related to activity-dependent plasticity in cortex, i.e., sleep spindles, slow oscillations, and delta waves^18,35–37^, and examined whether activity in DLS was also modulated during these rhythms (**Figure 4d**). We found that both LFP signals and spiking in DLS were significantly modulated during slow oscillations, delta waves, and sleep spindles detected in M1 (**Figure 4e&f; Supplemental Figure 4**). To compare corticostriatal transmission strength during these rhythms, we measured the short-latency cross correlation magnitude for the population of putatively connected M1 and DLS pairs using the spiking activity during each sleep rhythm. Importantly, we applied a previously established normalization method to isolate and subtract off the influence of firing rate changes or LFP phase-locking differences across NREM rhythms on cross correlations^38^. This revealed that sleep spindles were unique periods of boosted corticostriatal transmission strength, compared to slow oscillations or delta waves (**Figure 4g & Supplemental Figure 3c&d**). Altogether, this indicated that sleep spindles during NREM may be particularly relevant periods for activity-dependent plasticity within the corticostriatal network, given the high transmission of activity from M1 to DLS.

### Striatal reactivations during sleep spindles reflect cortical input

We next assessed whether sleep spindles, or other NREM rhythms, were significant predictors of day-to-day changes in behavioral consistency. We found that sleep spindle density (events/minute) during post-sleep, but not pre-sleep, was a significant predictor of day-to-day changes in reaching consistency (pre-sleep sleep spindles: r = 0.07, *P* = 0.70, post-sleep sleep spindles: r = 0.38, *P* = 0.01, Pearson’s *r*). Neither delta waves nor slow oscillations were significantly predictive of day-to-day changes in reaching consistency (pre-sleep delta waves: r = 0.02, *P* = 0.87, post-sleep delta waves: r = 0.03, *P* = 0.83, pre-sleep slow oscillations: r = −0.11, *P* = 0.51, post-sleep slow oscillations: r = −0.08, *P* = 0.60, Pearson’s *r*). Given the evidence that corticostriatal transmission is boosted during sleep spindles, a possible explanation for the unique relationship between post-training sleep spindle density and day-to-day increases in behavioral consistency is that sleep spindles drive activity-dependent corticostriatal plasticity that impacts behavior. If this were the case, we would expect relevant M1 and DLS neural populations to be preferentially engaged during sleep spindles after training. In fact, we found that reach modulated (RM) M1 units, characterized by a significant modulation of activity during the reaching action, were significantly more modulated during sleep spindles after training, while non-RM M1 units did not significantly change in modulation from pre-to post-sleep (**Figure 5a&b**; RM M1 units: *P* = 0.02, non-RM M1 units: *P* = 0.75, two-sample Kolmogorov–Smirnov test between distributions from pre- and post-sleep, followed by a shift test to assess how quartiles of the distributions differed; *P* values for the rest of Figure 3 reflect these statistical tests).

**Figure 5.**
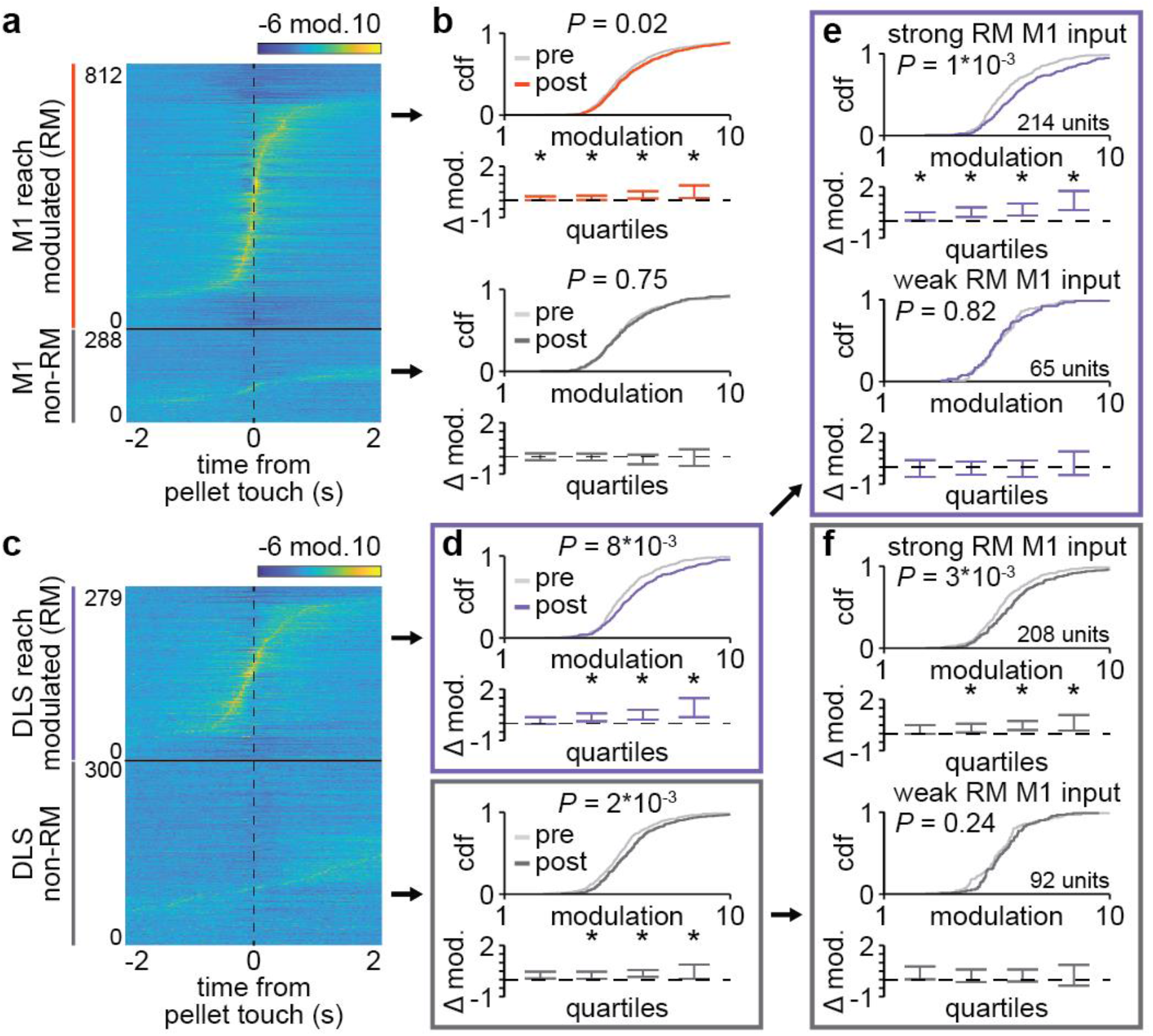
Striatal reactivations during sleep spindles reflect cortical input. **a.** Trial-averaged spiking activity during reaching for all reach modulated (RM) and non-RM M1 units across days and animals. **b.** Comparison of distributions of sleep spindle modulation during pre- and post-sleep for RM (top) and non-RM (bottom) M1 units, showing increased modulation from pre-to post-sleep specifically in RM M1 units. **c.** Trial-averaged spiking activity during reaching for all reach modulated (RM) and non-RM DLS units across days and animals. **d.** Comparison of distributions of sleep spindle modulation during pre- and post-sleep for RM (top) and non-RM (bottom) DLS units, showing increased modulation from pre-to post-sleep in both RM and non-RM DLS units. **e.** Comparison of distributions of sleep spindle modulation during pre- and post-sleep for RM DLS units with strong RM M1 input (top) and weak or no RM M1 input (bottom), showing increased modulation from pre-to post-sleep specifically in RM DLS units with strong RM M1 input. **f.** Comparison of distributions of sleep spindle modulation during pre- and post-sleep for non-RM DLS units with strong RM M1 input (top) and weak or no RM M1 input (bottom), showing increased modulation from pre-to post-sleep specifically in non-RM DLS units with strong RM M1 input.

How does this then affect downstream neural activity during sleep spindles in DLS? Surprisingly, both RM and non-RM DLS unit populations were significantly more modulated during sleep spindles after training (**Figure 5c&d**; RM DLS units: *P* = 8×10^-3^, non-RM DLS units: *P* = 2×10^-3^). One possibility is that this occurs because DLS activity during sleep spindles is driven by M1 input, rather than strictly reflecting reach modulation, as is the case for M1 neurons. Consistent with this, we were able to separate DLS unit populations that increased or did not increase in sleep spindle modulation after training based on putative connectivity with M1 units (**Figure 5e&f**; RM DLS units with strong RM M1 input: *P* = 1×10^-3^, RM DLS units with weak or no RM M1 input: *P* = 0.82, non-RM DLS units with strong RM M1 input: *P* = 3×10^-3^, non-RM DLS units with weak or no RM M1 input: *P* = 0.24; strong RM M1 input was defined as putative connectivity with three or more RM M1 units). In contrast, DLS units did not increase in modulation during either delta waves (**Supplemental Figure 5**) or slow oscillations (**Supplemental Figure 6**) after training. Altogether, this suggests that, while reactivations during sleep spindles reflect task modulation in M1, DLS reactivations reflect cortical input, suggesting a potential role for sleep spindles in reinforcing task-related corticostriatal connectivity, including novel connectivity such as projections from RM M1 units to previously non-RM DLS units.

### Sleep spindle modulation predicts offline changes in corticostriatal transmission strength

We next sought to directly examine whether sleep spindle modulation following training was related to modifications of the corticostriatal network within offline periods. To measure modifications of the corticostriatal network during offline periods, we calculated cross correlations of spiking activity across individual pairs of M1 and DLS units during the first and second half of each pre- and post-sleep period (**Figure 6a-c**). Cross correlations were generated specifically with spiking activity during NREM to control for any differences in time spent in each behavioral state. Consistent changes in the short-latency cross correlation magnitude from the first to second half of pre- or post-sleep would indicate a modification of corticostriatal transmission strength within the offline period. Given the evidence of reactivation during sleep spindles between RM M1 and putatively connected DLS units, we first specifically examined transmission strength changes within the pairs in this population that were significantly modulated to spindles (708/3,969 pairs in pre-sleep and 1,062/3,969 pairs in post-sleep). Strikingly, we observed an increase in corticostriatal transmission strength during post-sleep, but no significant change during pre-sleep (**Figure 6d&e**; pre-sleep: one-sample t-test: *t*(707) = −0.9, *P* = 0.37; post-sleep: one-sample t-test: *t*(1061) = 10.8, *P* = 6×10^-26^). Furthermore, increases in corticostriatal transmission strength across individual pairs of M1 and DLS units during post-sleep, but not pre-sleep, were correlated to the mean sleep spindle modulation of that pair (**Figure 6f&g**). Importantly, the amount of time spent in NREM was similar during pre- and post-sleep (**Supplemental Figure 7**). In contrast, transmission strength across RM M1 and putatively connected DLS unit pairs that were not significantly modulated to sleep spindles (1,164/3,969 pairs in pre-sleep and 989/3,967 pairs in post-sleep) did not change during either pre- or post-sleep and, within this population, changes in corticostriatal transmission strength across individual pairs of M1 and DLS units were not significantly correlated to the mean sleep spindle modulation of that pair (**Supplemental Figure 8**). These results suggested that, following training, offline corticostriatal transmission strength changes are linked to sleep spindle modulation.

**Figure 6.**
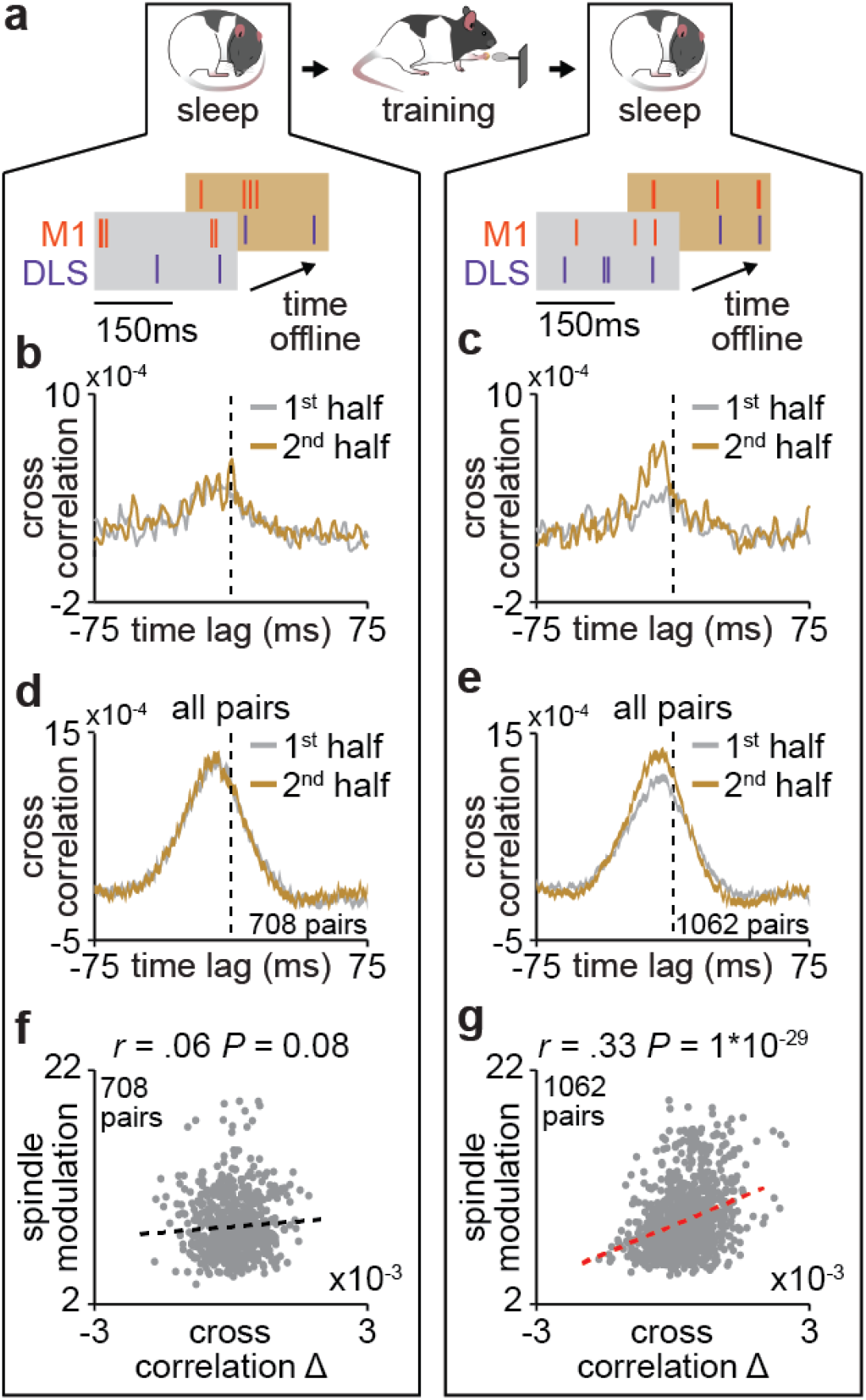
Sleep spindle modulation predicts offline changes in corticostriatal coupling. **a.** Schematic of NREM spiking activity snippets from example M1 and DLS units depicting the evolution of M1 and DLS spiking relationships from the first to second half of pre- (left) and post-sleep (right). **b.** Cross correlations of spiking activity during NREM from example M1 and DLS unit pair during the first and second half of pre-sleep, showing no change in cross correlation magnitude. **c.** Same as **b** for post-sleep, showing an increase in short-latency cross correlation magnitude. **d.** Cross correlations of spiking activity during NREM for all pairs of RM M1 and putatively connected DLS units that are significantly modulated to sleep spindles during the first and second half of pre-sleep, showing no change in cross correlation magnitude (width of line represents mean ± SEM). **e.** Same as **d** for post-sleep, showing an increase in short-latency cross correlation magnitude. **f.** Correlation between change in short-latency cross correlation magnitude and mean sleep spindle modulation for all pairs of RM M1 and putatively connected DLS units that are significantly modulated to sleep spindles during pre-sleep. **g.** Same as **f** for post-sleep.

### The interaction between sleep spindles and slow oscillations impact the role of sleep spindles within the corticostriatal network

To understand why changes in corticostriatal transmission strength occurred specifically in post-sleep, but not pre-sleep, we examined the interaction between sleep spindles and slow oscillations, a relationship known to be relevant for sleep-dependent processing^18,38,39^. We found that the distribution of temporal proximity to preceding slow oscillations in post-sleep significantly differed from the distribution in pre-sleep, with slow oscillations in closer proximity to sleep spindles during post-sleep (**Figure 7a**; *P*=2×10^-29^, two-sample Kolmogorov–Smirnov test). We found that this close proximity of slow oscillations to sleep spindles increased firing rates during sleep spindles in both M1 and DLS (**Figure 7b&c**, M1 units: *P* = 0.02, DLS units: *P* = 0.02, two-sample Kolmogorov–Smirnov tests, followed by a shift test to assess how quartiles of the distribution differed). Notably, the rate of sleep spindles within 500ms after a slow oscillation was correlated to the mean change in corticostriatal transmission strength across all pairs of M1 and DLS units during post-sleep (**Figure 7d**). This suggested that proximity to slow oscillations may be an important factor in whether sleep spindles drive plasticity. Altogether, our results provide evidence for offline plasticity within the corticostriatal network after training and that sleep spindles, and their interactions with slow oscillations, are important mediators of such plasticity.

**Figure 7.**
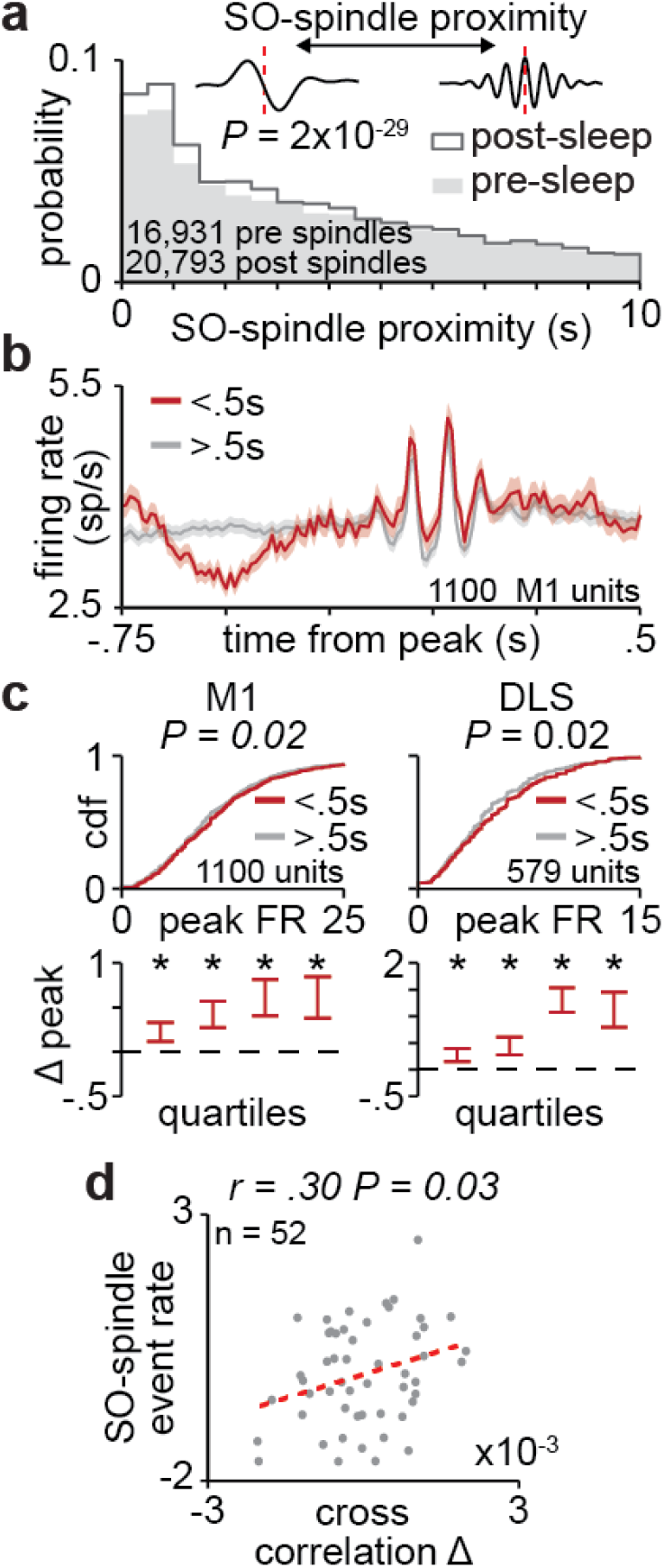
The interaction between sleep spindles and slow oscillations impact the role of sleep spindles within the corticostriatal network. **a.** Distributions of the temporal proximity to preceding slow oscillations for all sleep spindles during pre- and post-sleep across days and animals. **b.** Firing rate across M1 units during sleep spindles with close proximity to slow oscillations (<0.5 seconds) and all other sleep spindles (width of line represents mean ± SEM). **c.** Comparison of distributions of peak firing rates for M1 (left) and DLS (right) units during sleep spindles with close proximity to slow oscillations (<0.5 seconds) and all other sleep spindles. **d.** Correlation between each days’ post-sleep density of sleep spindles in close proximity to slow oscillations (<0.5 seconds; normalized by subtracting the rate in pre-sleep and normalized within each animal by z-scoring across days) and mean change in short-latency cross correlation across all pairs of RM M1 and putatively connected DLS units that are significantly modulated to sleep spindles (normalized by subtracting the change in pre-sleep and normalized within each animal by z-scoring across days).

## Discussion

Skilled behaviors exist in a continuum between being flexible - adapting quickly to changes in the environment - and automatic - inflexible to changes but cognitively efficient^1–4^. Plasticity within the corticostriatal network is thought to regulate the balance between flexibility and automaticity^4,5,26^. In this study, we study long-term training that resulted in automaticity, as evidenced by day-to-day invariance in reaching behavior that persisted even when the food pellet was moved such that reaches were no longer successful in retrieving the pellet. We show that, during such long-term training, corticostriatal functional connectivity increased during offline periods and provide evidence that sleep spindles uniquely engage the corticostriatal network to mediate such plasticity.

Our results provide evidence that sleep plays an important role in modifying cross-area connectivity during learning. While coordinated cross-area reactivations during sleep have been reported across several brain networks^19,40–45^, how such coordinated activity patterns precisely shape the connectivity across brain regions remains largely unexplored (but see ^19^). For example, studies informing the *systems consolidation theory* have posited that coordinated activity patterns across the hippocampus and cortex during sleep drive intra-cortical plasticity^23,25,46^. However, it is not known whether sleep impacts hippocampal-cortical connectivity, despite evidence for a change in hippocampal-cortical coupling after learning^47^. Our results thus suggest that the *systems consolidation theory* may need to be broadened to consider a role for sleep in increasing the coupling between connected regions that can impact subsequent wake network activity and behavior. As evidence for cross-area reactivations in different brain networks continues to grow^48^, it will be important to consider how such activity patterns may impact both local and cross-area plasticity.

Our measures of cross-area connectivity are based on the coordination of LFP signals and single unit spike timing across M1 and DLS. We observed a subset of both LFP electrodes and corticostriatal neuron pairs showing evidence of increased connectivity with training, indicating the selective strengthening of corticostriatal connectivity - but what is the neural basis for these changes? One possibility is that our functional measures of connectivity reflect changes in synaptic strength of M1 projections to the DLS. This is consistent with evidence for the strengthening of cortical inputs to the striatum with motor training^49^. An alternative possibility is that coordinated inputs to both M1 and DLS drive increased functional connectivity. We believe our results are most consistent with a physical change in synaptic strength, as we observed evidence of increased cross-area connectivity in two distinct states, NREM, reflected as increased LFP coherence, and awake task performance, reflected in the emergence of predictable cross-area dynamics. Future work is required to determine whether our observations are consistent with structural changes in synaptic strength.

We also provide evidence that sleep spindles are uniquely poised to mediate the enhancement of corticostriatal coupling during the offline period following training. While sleep spindles have been previously suggested to be important for plasticity^36,50^, the precise link between sleep spindles, plasticity, and behavior has remained unclear. Here we also show that striatal NMDA activation during the offline periods following training is required for increases in behavioral consistency. This suggests that sleep spindles may be important drivers of corticostriatal plasticity through NMDA activation. This is consistent with work showing that corticostriatal plasticity is NMDA-dependent^13,14^, as well as *in vitro* work examining how sleep spindle activity patterns might drive plasticity^50^. Additionally, we provide evidence that the proximity of sleep spindles to preceding slow oscillations is an important regulator of plasticity, consistent with previous work^18,39^. As slow oscillations have been linked to NMDA receptor activation^51^, one intriguing possibility is that slow oscillations gate sleep spindle plasticity through the activation of NMDA receptors.

Our results link offline corticostriatal plasticity to the emergence of predictable low-dimensional cross-area activity. It has been previously demonstrated that M1 exhibits consistent low-dimensional population neural dynamics during consistently produced motor actions^52^. There is also growing evidence that subcortical regions such as the DLS are important for stabilizing cortical activity patterns and the emergence of consistent behaviors^8,12^. Consistent with this idea, task-related coordination of M1 and DLS activity emerges with skill acquisition^8–10^. Here we link offline increases in corticostriatal functional connectivity to the ability to predict low-dimensional population activity in DLS from M1 activity. This suggests a model in which consistent low-dimensional neural dynamics emerge across the motor network with training and that motor network plasticity during offline periods is important for the emergence of such consistent cross-area dynamics.

Lastly, we link offline corticostriatal plasticity in the corticostriatal network to the emergence of fast and consistent reaching behavior, reflected in the invariance of day-to-day reaching velocity profile. This is consistent with a range of studies demonstrating that sleep benefits speed and consistency in motor tasks in humans^53,54^ and rodents^37,55^, as well as rodent brain-machine interface (BMI) tasks^17,18^. Therefore, our results suggest the possibility that a fundamental role of sleep is to modify the corticostriatal network to impact the consistency of behavior in a range of tasks. Further work is required to determine the precise role of offline corticostriatal plasticity in different contexts. One important avenue of research is to explore whether sleep can impact corticostriatal connectivity in the context of maladaptive automatic behaviors, such as addiction, that have been linked to the corticostriatal network^26,56^. Notably, there is evidence that the reactivation of a stored memory can make the memory temporarily labile^27^ and recent work has shown that the modulation of NREM rhythms can regulate modulate memory consolidation vs. forgetting^18^. Therefore, it will be informative to determine whether similar manipulations could be used in the context of maladaptive automatic behaviors to provide a therapeutic benefit.

---

**Supplemental Figure 1.**
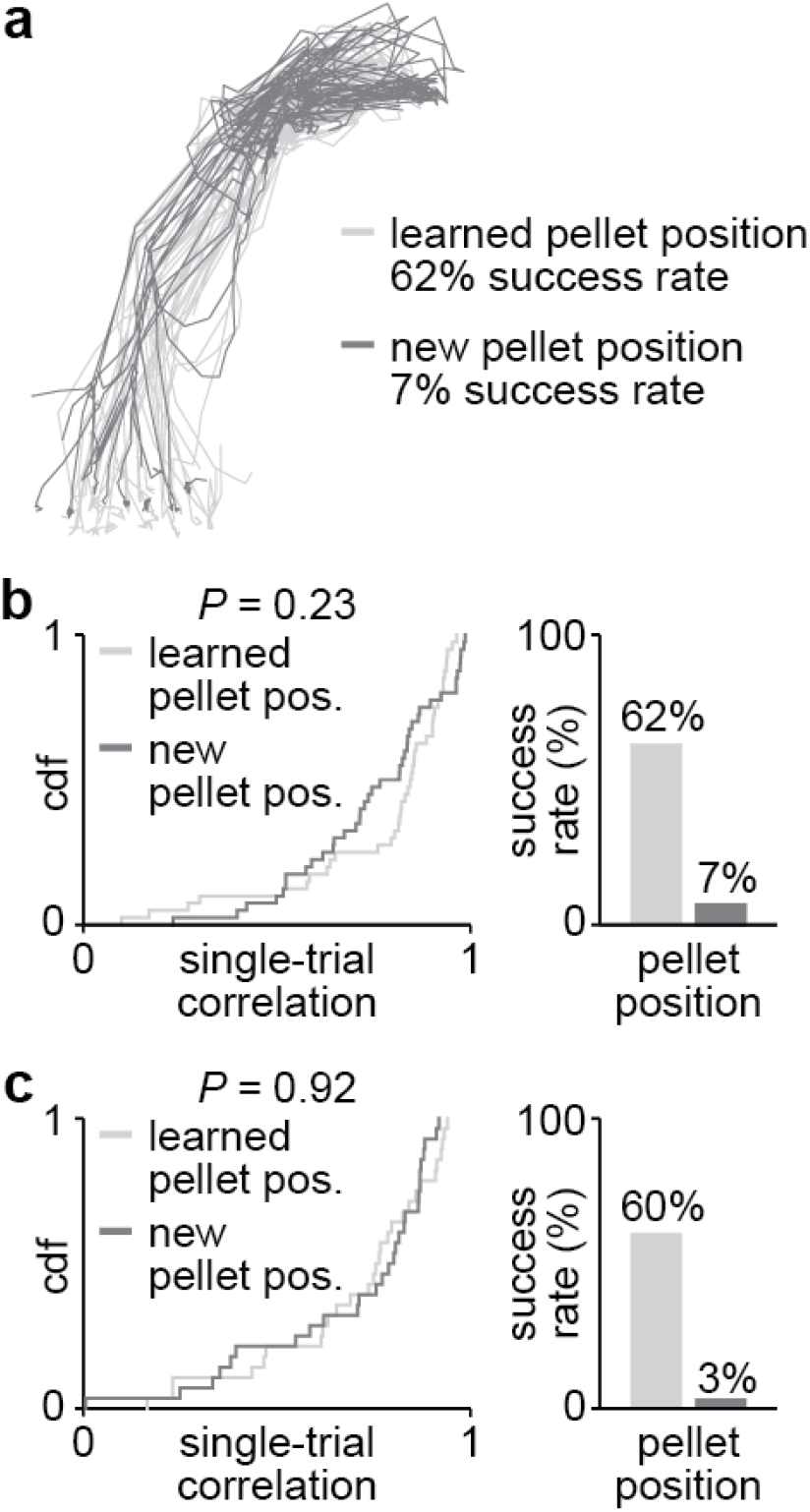
Animals do not adapt quickly to new pellet position after long-term reach-to-grasp training. **a.** Reach trajectories from example session following reach-to-grasp task training paradigm, showing that reach trajectories with the pellet in the learned position or a new position are largely overlapping. **b.** Histogram of single-trial correlation values for individual reach trajectories to the mean reach trajectory for trials with the pellet in the learned pellet position or a new pellet position in first example animal (left; two-sample Kolmogorov–Smirnov test between distributions from trials with pellet in learned position and pellet in new position) and success rate in pellet retrieval for learned and new pellet positions (right). **c.** same as **b** for second example animal.

**Supplemental Figure 2.**
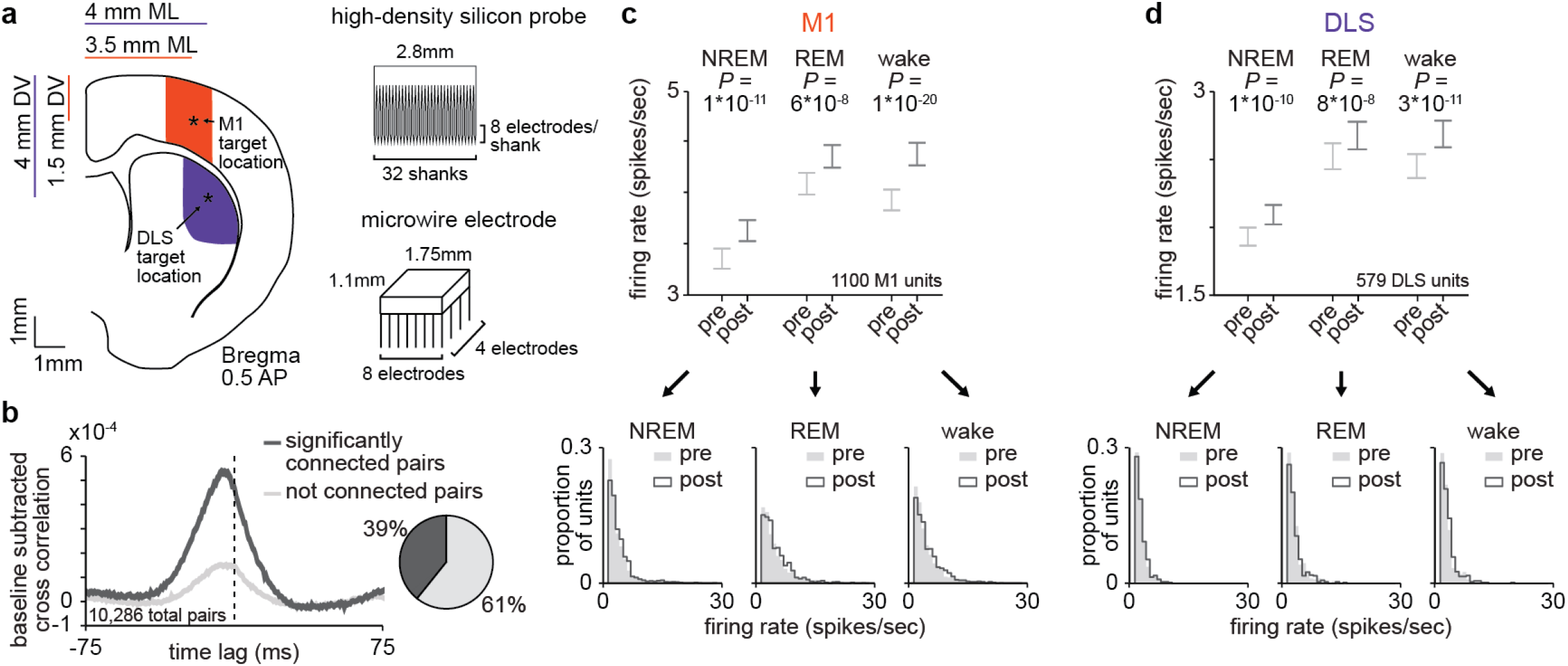
Electrophysiology recordings from M1 and DLS. **a.** Illustration of target electrode locations in M1 and DLS and dimensions of microwire electrode and silicon probes. **b.** Cross correlations of spiking activity across all pairs of putatively connected and not putatively connected pairs of M1 and DLS units (width of line represents mean ± SEM) and pie chart depicting percentage of all pairs that are classified as significantly connected. **c.** Comparison of firing rates in M1 across behavioral states, before and after training (1,100 M1 units; 3.4 ± 0.1 spikes/second in pre NREM vs. 3.7 ± 0.1 spikes/second in post NREM, mean ± SEM, paired-sample t-test: *t*(1099) = −6.8, *P* = 1×10^-11^; 4.2 ± 0.1 spikes/second in pre REM vs. 4.4 ± 0.1 spikes/second in post REM, mean ± SEM, paired-sample t-test: *t*(1099) = −5.5, *P* = 6×10^-8^; 4.0 ± 0.1 spikes/second in pre wake vs. 4.4 ± 0.1 spikes/second in post wake, mean ± SEM, paired-sample t-test: *t*(1099) = −9.5, *P* = 1×10^-20^). **d.** Comparison of firing rates in DLS across behavioral states, before and after training (579 DLS units; 1.9 ± 0.1 spikes/second in pre NREM vs. 2.1 ± 0.1 spikes/second in post NREM, mean ± SEM, paired-sample t-test: *t*(578) = −6.5, *P* = 1×10^-10^; 2.5 ± 0.1 spikes/second in pre REM vs. 2.7 ± 0.1 spikes/second in post REM, mean ± SEM, paired-sample t-test: *t*(578) = −5.5, *P* = 8×10^-8^; 2.4 ± 0.1 spikes/second in pre wake vs. 2.8 ± 0.1 spikes/second in post wake, mean ± SEM, paired-sample t-test: *t*(578) = −6.8, *P* = 3×10^-11^).

**Supplemental Figure 3.**
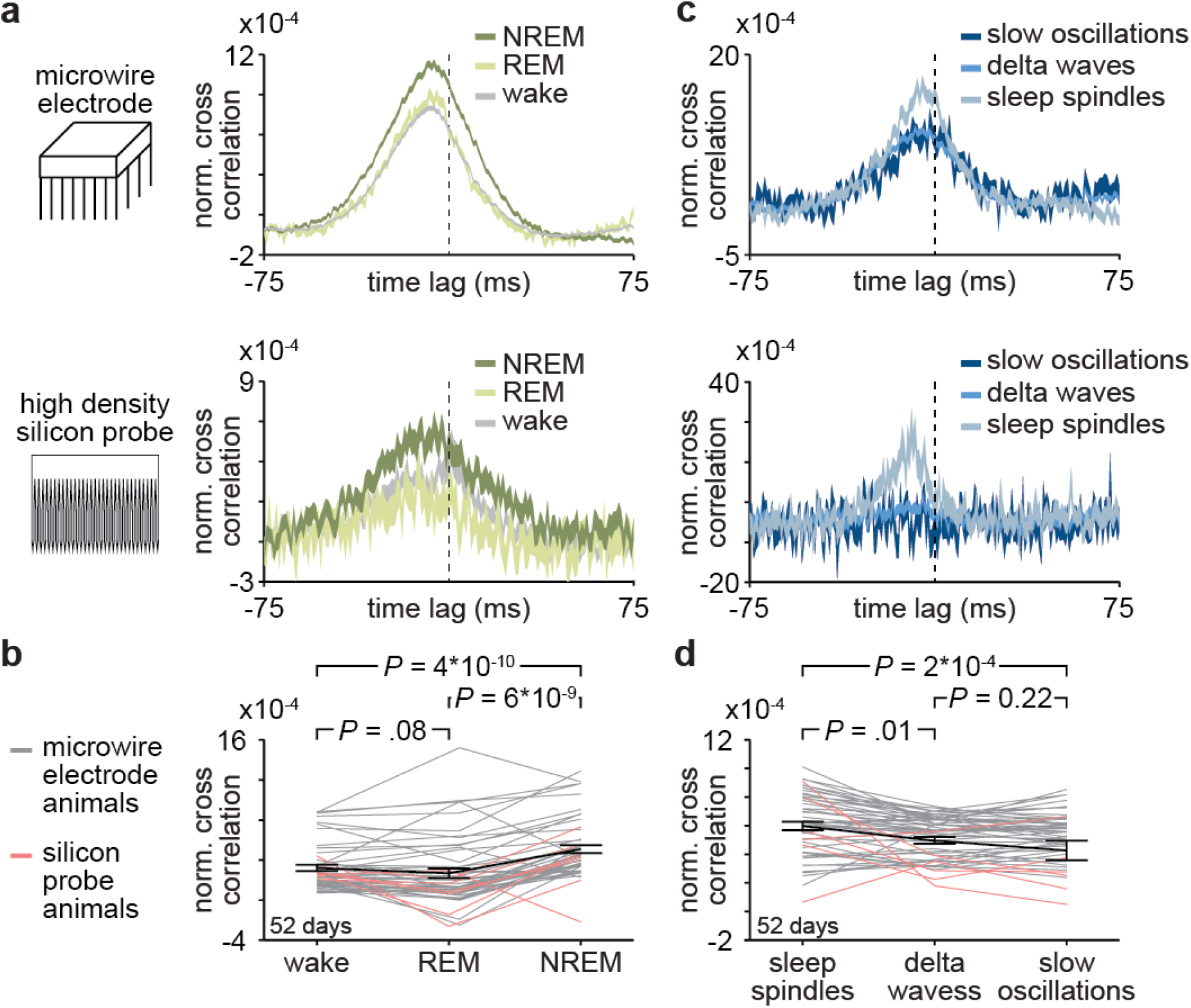
Corticostriatal transmission strength across behavioral states and NREM rhythms. **a.** Comparison of normalized cross correlations across behavioral states for all putatively connected pairs of M1 and DLS units in example animals implanted with either microwire electrode array (top) or high-density silicon probe (bottom), showing that corticostriatal transmission strength is maximal in NREM sleep (width of line represents mean ± SEM) **b.** Comparison of mean short-latency correlation magnitude (1-10ms time lag) across behavioral states, values from days for animals implanted with microwires in grey, silicon probes in red, and mean ± SEM across animals in black (*n* = 52 days across 7 rats; wake: 3.2*10^-4^ ± 0.3*10^-4^ correlation value, REM: 2.7*10^-4^ ± 0.5*10^-4^ correlation value, NREM: 5.1*10^-4^ ± 0.4*10^-4^ correlation value; wake vs. REM: *t*(51) = 1.8, *P* = 0.08, paired-sample t-test, REM vs. NREM: *t*(51) = −7.0, *P* = 6*10^-9^, paired-sample t-test, wake vs. NREM: *t*(51) = −7.7, *P* = 4×10^-10^, paired-sample t-test). **c.** Comparison of normalized cross correlations across NREM rhythms for all putatively connected pairs of M1 and DLS units in example animals implanted with either microwire electrode array (top) or high-density silicon probe (bottom), showing that corticostriatal transmission strength is maximal during sleep spindles (width of line represents mean ± SEM). **d.** Comparison of mean short-latency correlation magnitude (1-10ms time lag) across NREM rhythms, values from days for animals implanted with microwires in grey, silicon probes in red, and mean ± SEM across animals in black (*n* = 52 days across 7 rats; sleep spindles: 9.9*10^-4^ ± 0.8*10^-4^ correlation value, delta waves: 7.5*10^-4^ ± 0.6*10^-4^ correlation value, slow oscillations: 5.7*10^-4^ ± 1.7*10^-4^ correlation value; sleep spindles vs. slow oscillation: *t*(51) = 4.0, *P* = 2*10^-4^, paired-sample t-test, delta waves vs. slow oscillations: *t*(51) = 1.2, *P* = 0.22, paired-sample t-test, sleep spindles vs. delta waves: *t*(51) = 2.4, *P* = 0.01, paired-sample t-test).

**Supplemental Figure 4.**
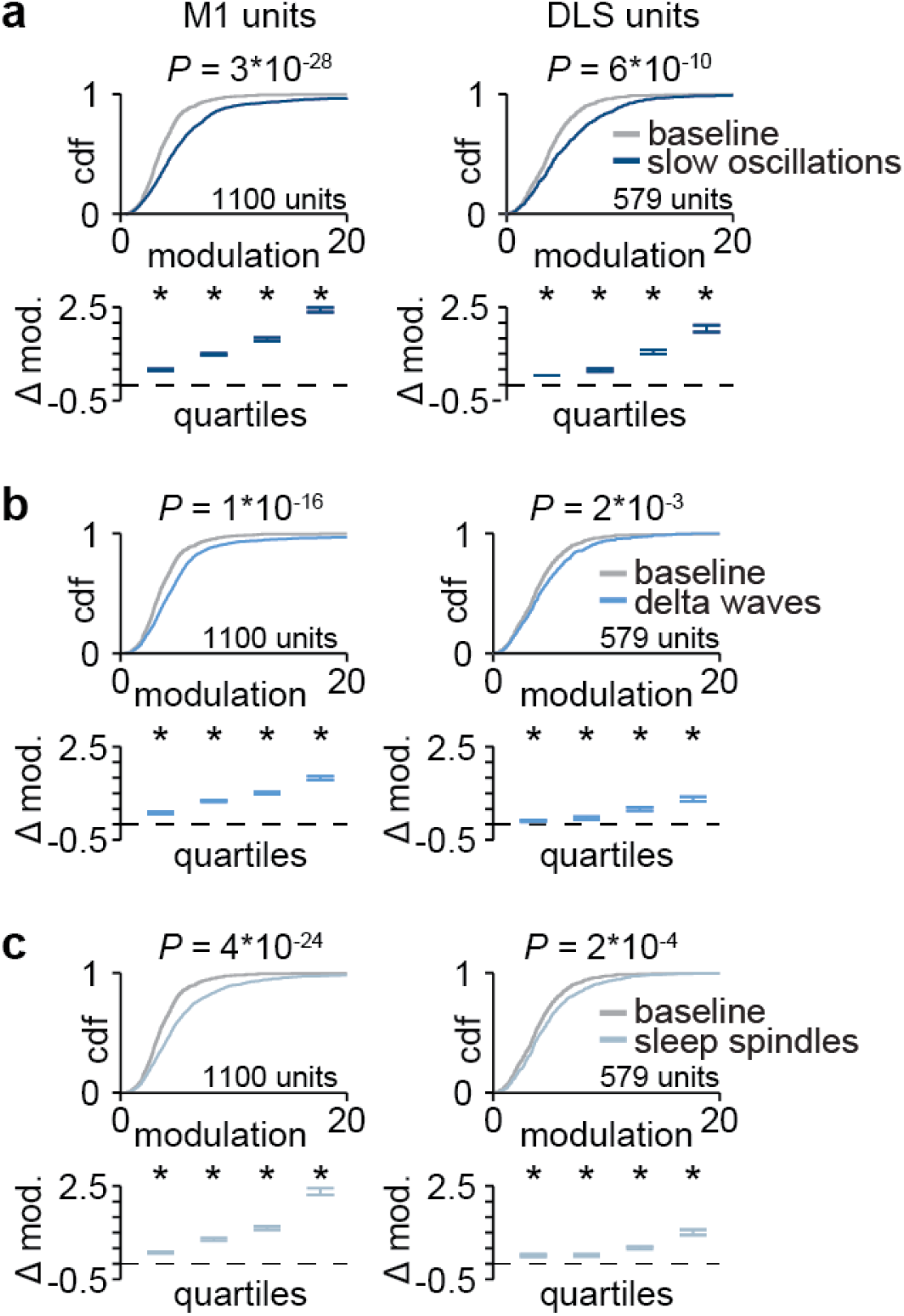
Corticostriatal modulation across NREM rhythms. **a.** Comparison of firing rate modulation distributions for M1 (left) and DLS (right) units during slow oscillations and a baseline NREM period (M1: *P* = 3*10^-28^, DLS: *P* = 6*10^-10^, two-sample Kolmogorov– Smirnov test, followed by a shift test to assess how quartiles of the distribution differed). **b.** Comparison of firing rate modulation distributions for M1 (left) and DLS (right) units during delta waves and a baseline NREM period (M1: *P* = 1 *10^-16^, DLS: *P* = 2*10^-3^, two-sample Kolmogorov– Smirnov test, followed by a shift test to assess how quartiles of the distribution differed). **c.** Comparison of firing rate modulation distributions for M1 (left) and DLS (right) units during sleep spindles and a baseline NREM period (M1: *P* = 4*10^-24^, DLS: *P* = 2*10^-4^, two-sample Kolmogorov–Smirnov test, followed by a shift test to assess how quartiles of the distribution differed).

**Supplemental Figure 5.**
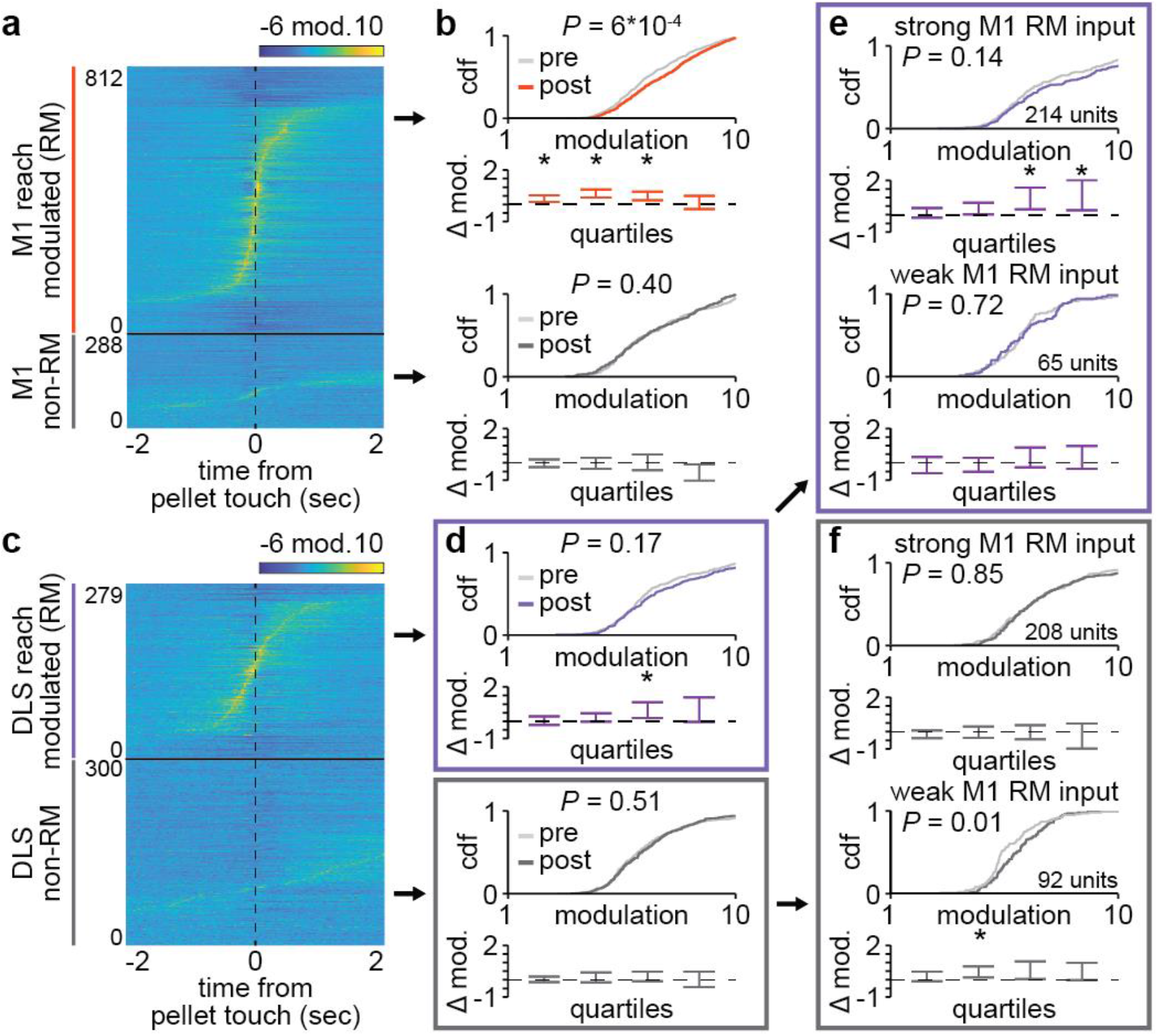
Delta wave modulation change with training. **a.** Trial-averaged spiking activity during reaching for all reach modulated (RM) and non-RM M1 units across days and animals. **b.** Comparison of distributions of delta wave modulation during pre- and post-sleep for RM (top) and non-RM (bottom) M1 units (RM M1 units: *P* = 6×10^-4^, non-RM M1 units: *P* = 0.40, two-sample Kolmogorov–Smirnov test between distributions from pre- and post-sleep, followed by a shift test to assess how quartiles of the distributions differed; *P* values for the rest of Supplemental Figure 5 legend reflect these statistical tests). **c.** Trial-averaged spiking activity during reaching for all reach modulated (RM) and non-RM DLS units across days and animals. **d.** Comparison of distributions of delta wave modulation during pre- and post-sleep for RM (top) and non-RM (bottom) DLS units (RM DLS units: *P* = 0.17, non-RM M1 units: *P* = 0.51). **e.** Comparison of distributions of delta wave modulation during pre- and post-sleep for RM DLS units with strong RM M1 input (top) and weak or no RM M1 input (bottom; RM DLS units with strong RM M1 input: *P* = 0.14, RM DLS units with weak or no RM M1 input: *P* = 0.72). **f.** Comparison of distributions of delta wave modulation during pre- and post-sleep for non-RM DLS units with strong RM M1 input (top) and weak or no RM M1 input (bottom; non-RM DLS units with strong RM M1 input: *P* = 0.85, non-RM DLS units with weak or no RM M1 input: *P* = 0.01).

**Supplemental Figure 6.**
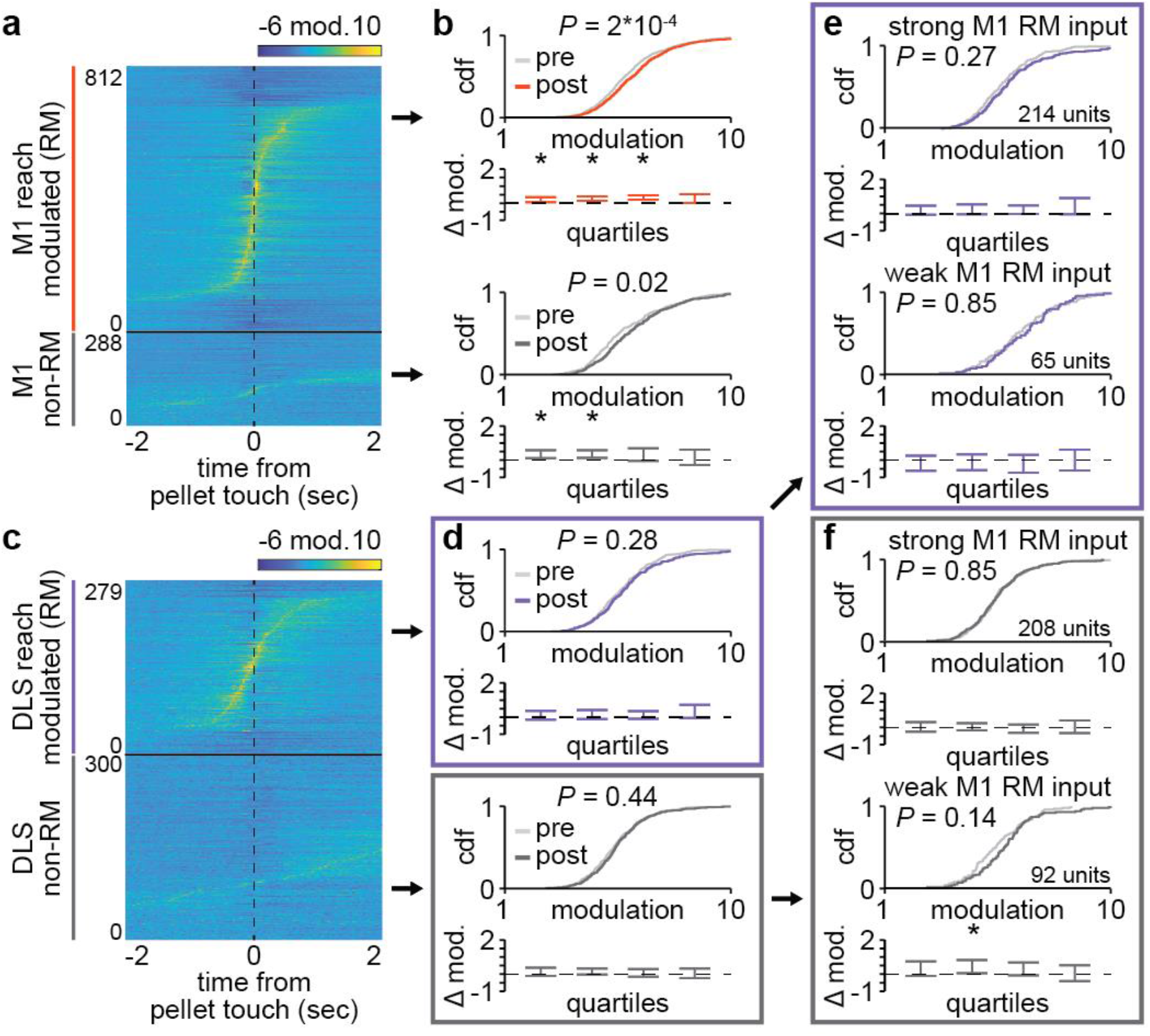
Slow oscillation modulation change with training. **a.** Trial-averaged spiking activity during reaching for all reach modulated (RM) and non-RM M1 units across days and animals. **b.** Comparison of distributions of slow oscillation modulation during pre- and post-sleep for RM (top) and non-RM (bottom) M1 units (RM M1 units: *P* = 2×10^-4^, non-RM M1 units: *P* = 0.02, two-sample Kolmogorov–Smirnov test between distributions from pre- and post-sleep, followed by a shift test to assess how quartiles of the distributions differed; *P* values for the rest of Supplemental Figure 6 legend reflect these statistical tests). **c.** Trial-averaged spiking activity during reaching for all reach modulated (RM) and non-RM DLS units across days and animals. **d.** Comparison of distributions of slow oscillation modulation during pre- and post-sleep for RM (top) and non-RM (bottom) DLS units (RM DLS units: *P* = 0.28, non-RM M1 units: *P* = 0.44). **e.** Comparison of distributions of slow oscillation modulation during pre- and post-sleep for RM DLS units with strong RM M1 input (top) and weak or no RM M1 input (bottom; RM DLS units with strong RM M1 input: *P* = 0.27, RM DLS units with weak or no RM M1 input: *P* = 0.85). **f.** Comparison of distributions of slow oscillation modulation during pre- and post-sleep for non-RM DLS units with strong RM M1 input (top) and weak or no RM M1 input (bottom; non-RM DLS units with strong RM M1 input: *P* = 0.85, non-RM DLS units with weak or no RM M1 input: *P* = 0.14).

**Supplemental Figure 7.**
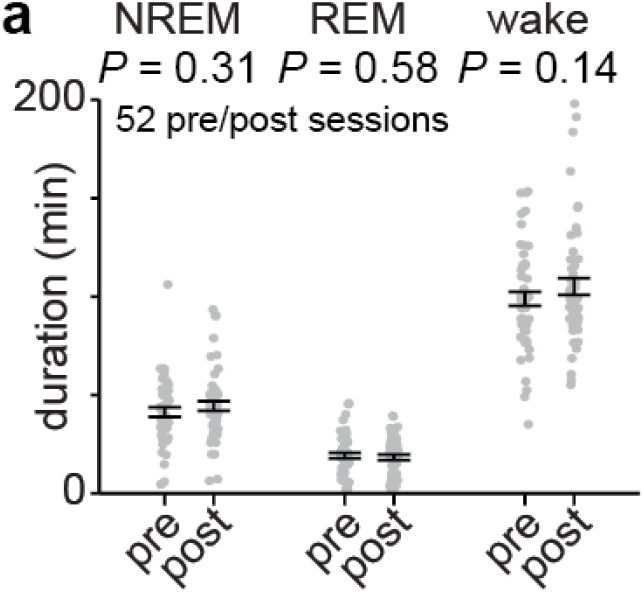
Comparison of time spent in each behavioral state during pre- and post-sleep. **a.** Comparison of durations spent in each behavioral state during pre- and post-sleep across days and animals (NREM: 41.2 ± 2.4 minutes of pre NREM vs. 44.3 ± 2.5 minutes of post NREM, mean ± SEM, paired-sample t-test: *t*(51) = −1.03, *P* = 0.31; REM: 17.2 ± 1.4 minutes of pre REM vs. 16.3 ± 1.4 minutes of post REM, mean ± SEM, paired-sample t-test: *t*(51) = 0.56, *P* = 0.58; wake: 98.8 ± 3.6 minutes of pre wake vs. 106.0 ± 4.3 minutes of post wake, mean ± SEM, paired-sample t-test: *t*(51) = −1.5, *P* = 0.14).

**Supplemental Figure 8.**
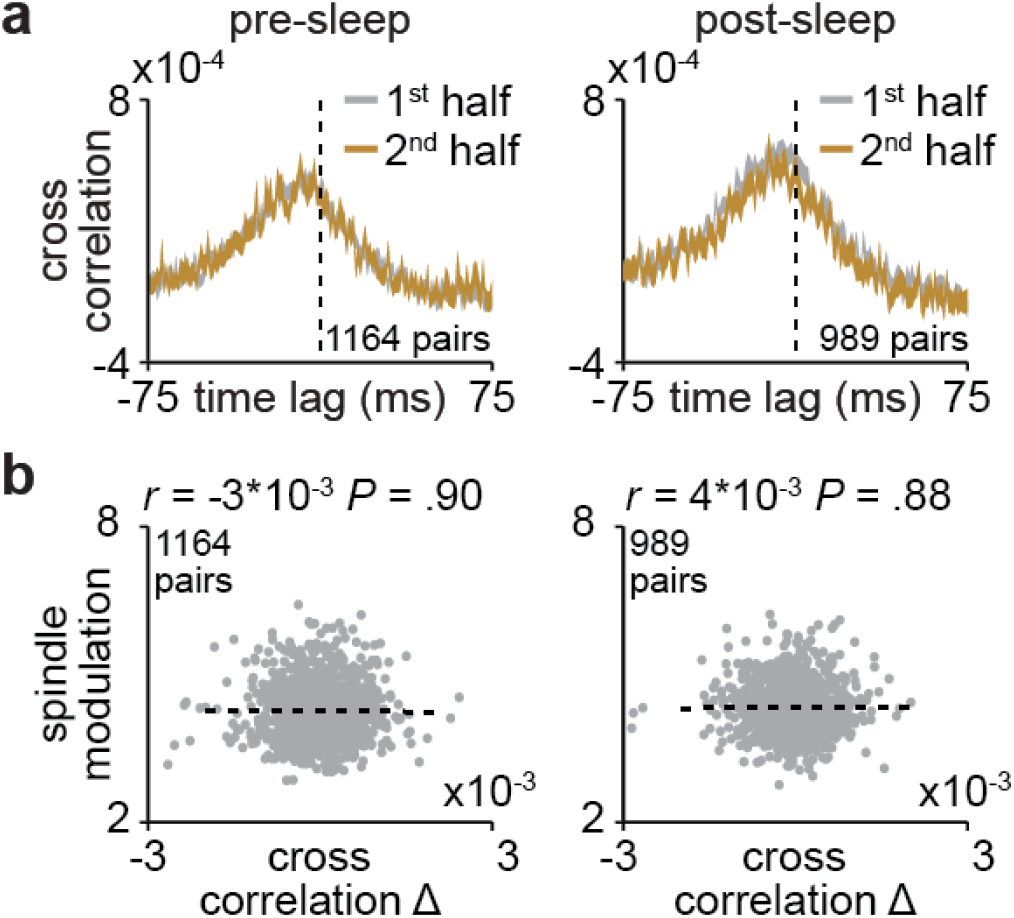
Corticostriatal transmission strength changes for non-sleep spindle modulated pairs of M1 and DLS units. **a.** Cross correlations of spiking activity during NREM for all pairs of RM M1 and putatively connected DLS units that are not significantly modulated to sleep spindles during the first and second half of pre-sleep (left) and post-sleep (right), showing no changes in cross correlation magnitude (width of line represents mean ± SEM) **b.** Correlation between change in short-latency cross correlation magnitude and mean sleep spindle modulation for all pairs of RM M1 and putatively connected DLS units that are not significantly modulated to sleep spindles during pre-sleep (left) and post-sleep (right).

## Methods

### Animal care and surgery (Supplemental Figure 2)

This study was performed in strict accordance with guidelines from the USDA Animal Welfare Act and United States Public Health Science Policy. Procedures were in accordance with protocols approved by the Institutional Animal Care and Use Committee at the San Francisco Veterans Affairs Medical Center. This study consists of experiments performed with fourteen male Long-Evans rats (approximately 12-16 weeks old), housed under controlled temperature and a 12-h light/12-h dark cycle with lights on at 6:00 a.m. Animal experiments were performed during the light period. All surgical procedures were performed using sterile techniques under 2–4% isoflurane. Six animals were implanted with either microwire electrodes (*n* = 4 animals; 32 or 64 channel 33μm diameter Tungsten microwire arrays with ZIF-clip adapter; Tucker-Davis Technology) or high-density silicon probes (*n* = 2 animals; 256 channel custom-built silicon probes) targeted to both the forelimb area of M1, centered at 3.5mm lateral and 0.5mm anterior to bregma and implanted in layer V at a depth of 1.5mm, and the DLS, centered at 4mm lateral and 0.5mm anterior to bregma and implanted at a depth of 4mm. Six additional animals were implanted with infusion cannulas (PlasticsOne; 26Ga) targeted to the DLS. Surgery involved exposure and cleaning of the skull, preparation of the skull surface (using cyanoacrylate), and implantation of skull screws for overall headstage stability. In the animals implanted with neural probes, a reference screw was implanted posterior to lambda, contralateral to the neural recordings and a ground screw was implanted posterior to lambda, ipsilateral to the neural recordings. Craniotomy and durectomy were then performed, followed by implantation of neural probes or infusion cannulas and securing of the implant with C&B Metabond (Parkell, Product #S380) and Duralay dental acrylic (Darby, Product #8830630). In four of the animals implanted with neural probes, the forearm was also implanted with a pair of twisted electromyography (EMG) wires (0.007” single-stranded, Teflon-coated, stainless steel wire; A-M Systems) with a hardened epoxy ball (J-B Weld Company) at one end preceded by 1–2mm of uncoated wire under the ball. Wires were inserted into the muscle belly and pulled through until the ball came to rest on the belly. EMG wires were braided, tunneled under the skin to a scalp incision and soldered into an electrode interface board (ZCA-EIB32; Tucker-Davis Technology). The postoperative recovery regimen included administration of buprenorphine at 0.02mg/kg and meloxicam at 0.2mg/kg. Dexamethasone at 0.5mg/kg and trimethoprim/sulfadiazine at 15mg/kg were also administered postoperatively for 5 days. All animals recovered for at least one week before the start of behavioral training.

### *In vivo* electrophysiology

Units, local field potentials (LFP), and EMG activity were recorded using an RZ2 system (Tucker-Davis Technologies). For the microwire animals, spike data was sampled at 24,414Hz and LFP/EMG data at 1,017Hz. To record spiking data in these animals, thresholds for spiking activity were set online using a standard deviation of 4.5 (calculated over a 1-min baseline period using the RZ2 system). Waveforms and timestamps were stored for any event that crossed that threshold. Spike sorting was then performed using Offline Sorter v.4.3.0 (Plexon) with a principal component analysis-based clustering method followed by manual inspection. Spikes were sorted separately for each day, combining pre-sleep, training, and post-sleep sessions. We accepted units based on waveform shape, clear cluster boundaries in principal component space and 99.5% of detected events with an ISI>2ms. For silicon probe animals, signals were recorded at 24,414Hz. In these animals, spike times and waveforms were detected from the broadband signal using Offline Sorter v.4.3.0 (Plexon). Spike waveforms were then sorted using Kilosort2 (https://github.com/MouseLand/Kilosort2). We accepted units based on manual inspection using Phy (https://github.com/cortex-lab/phy) and 99.5% of detected events with an ISI>2ms.

### Viral injection (Figure 1a)

To label anterograde projections in M1 we injected 750nl of AAV8-hsyn-JAWs-KGC-GFP-ER2 virus into two sites (1.5mm anterior, 2.7mm lateral to bregma, at a depth of 1.4mm and 0.5 posterior, 3.5mm lateral to bregma, at a depth of 1.4mm). Two weeks after injection rats were anesthetized and transcardially perfused with 0.9% sodium chloride, followed by 4% formaldehyde. The harvested brains were post-fixed for 24 h and immersed in 20% sucrose for 2 days. Coronal cryostat sections (40-μm thickness) were then mounted and imaged with a fluorescent microscope.

### Reach-to-grasp task (Figure 1; Supplemental Figure 1)

Rats naïve to any motor tasks were first tested for forelimb preference. This involved presenting approximately ten food pellets to the animal and observing which forelimb was most often used to reach for the pellet. Rats then underwent surgery for either neural probe or cannula implantation in the hemisphere contralateral to preferred paw. Following the one-week recovery period, rats were trained using an automated reach-box, controlled by custom MATLAB scripts and an Arduino microcontroller. This setup requires minimal user intervention, as described previously (Wong, et al., 2015). Each trial consisted of a pellet dispensed on the pellet tray followed by an alerting beep indicating that the trial was beginning, then the door would open. Animals had to reach, grasp, and retrieve the pellet. A real-time ‘pellet detector’ using an infrared sensor centered over the pellet was used to determine when the pellet was moved, indicating the trial was over and then the door was closed. All trials were captured by a camera placed on the side of the behavioral box (*n* = 2 animals monitored with a Microsoft LifeCam at 30 frames/second; *n* = 12 animals monitored with a Basler ace acA640-750uc at 75 frames/second). For animals implanted with neural probes, each animal underwent five to fourteen days of training (~100–150 trials per day). For the infusion cannula implanted animals, each animal underwent ten days of training (100 trials per day). Rats had fifteen seconds to complete each trial, and trials were separated by a ten second inter-trial-interval. Reach trajectories were captured from video using DeepLabCut (Mathis, et al., 2018) to track the center of the rat’s paw as well as the food pellet. Reach trajectories consisted of the paw trajectory from 500ms before to 500ms after “pellet touch”, which was classified as the frame in which the paw was closest to the pellet, before the pellet was displaced off the pellet holder. Only trials in which the pellet was displaced off the pellet holder were considered. We assessed behavioral consistency throughout training in both neural probe and cannula implanted animals by calculating the correlation between the mean velocity profile of reaches on each day of training and the mean velocity profile of reaches on the last day of training. These correlations were computed separately for the x and y dimensions and then averaged. At the end of training, we tested whether reaching behavior was automatic in two of the neural probe implanted animals by performing a 100 trial training session on the subsequent day with the pellet moved to a new location (~10mm lateral from original pellet position) and observing whether the animal’s reaching behavior changed. We performed a similar experiment for two additional animals naïve to the task and without neural implant to test whether reaching was flexible or automatic at the start of training. These animals performed ~200 trials on two consecutive days. To calculate single-trial reach trajectory correlations, we first generated a mean trajectory in each dimension (x and y) for trials with the pellet in the learned position and trials with the pellet in the new position (mean trajectories were computed separately for each pellet position). Single trial trajectories were then correlated to the mean trajectory in each dimension and then averaged across the x and y dimension. To compare across pellet positions, we considered reach trajectories up to pellet touch (from 500ms before pellet touch to pellet touch), as automatic reaches with the pellet in the new position often missed the pellet and pellet holder completely.

### DLS infusions (Figure 1)

To test if blocking the activation of striatal NMDA receptors during the offline period after training disrupts increases in behavioral consistency, we infused either 1ul of saline or NMDA blocker AP5 (5μg/μl) at an infusion rate 200nl/minute into the DLS immediately following training in six animals for ten consecutive days. In the first five days of training, we infused three rats with AP5 and three rats with saline, for the second five days, we switched the infusion, i.e., animals that received AP5 in the first five days, received saline for the second five days, and vice-versa.

### Sleep classification (Figure 4a; Supplemental Figure 7)

All neural data analyses were conducted using MATLAB 2019a (MathWorks) and functions from the EEGLAB (http://sccn.ucsd.edu/eeglab/) and Chronux (http://chronux.org/) toolboxes. Sleep was classified using cortical LFP signals and movement measured by video or EMG activity. LFP was preprocessed by artifact rejection, including manual rejection of noisy channels and z-scoring of each channel across the entire recording session. A mean LFP channel was then generated in M1 for sleep classification by averaging across all M1 channels. This mean M1 LFP channel was then segmented into non-overlapping 10 second windows. In each window the power spectral density was computed using the Chronux function *mtspecgramc* and then averaged over the delta (1 –4Hz) and theta (5-10Hz/2-15Hz) frequency bands. Both LFP power bands were then normalized by z-scoring. Epochs with high delta power (>0 z-scored delta) and no movement were classified as NREM, epochs with high theta and low delta power (>0 z-scored theta and <0 z-scored delta) were classified as REM sleep, and other epochs were classified as wake (Watson, et al., 2016). All consecutive NREM or REM epochs that were less than 30 seconds long (3 consecutive epochs) were reclassified as wake.

### Assessing corticostriatal functional connectivity using LFP coherence (Figure 2; Figure 3)

To measure corticostriatal functional connectivity across days, we measured LFP coherence during NREM across all M1 and DLS electrode pairs on each pre- and post-sleep session using chronux function *cohgramc*. For these analyses, we first applied common-mode referencing using the median signal, i.e., at every time-point, the median signal across all channels in a region was calculated and subtracted from every channel to decrease common noise and minimize volume conduction. Common-mode referencing was performed independently for the channels in each region, i.e., M1 and DLS. We classified “high coherence LFP pairs” as electrodes with a mean 4-8Hz coherence >0.6. To compare online changes in LFP coherence (from pre-to post-sleep on the same day) to offline changes in LFP coherence (from post-sleep on one day to pre-sleep on the next day), we computed a single value per pair for both online and offline coherence changes by averaging values across days of training.

### Predicting cross-area activity (Figure 3)

To assess cross-area dynamics, we first extracted low-dimensional representations of DLS activity by performing principal component analysis (PCA) on trial-averaged activity of DLS neurons time-locked to pellet touch and binned at 100ms, specifically for time bins from five seconds before to five seconds after pellet touch. Principal components were computed using MATLAB function *pca*. Spiking activity from five seconds before to five seconds after pellet touch and binned at 100ms was then projected onto each of the first two components to generate low-dimensional neural trajectory representations of population activity in DLS. We then fit a linear regression model to predict DLS reach-related neural trajectories from one second before to one second after pellet touch from single unit spiking activity in M1. A separate model was used to predict each principle component, using MATLAB function *fitlm* and five-fold cross validation. For each time bin of the neural trajectory, the preceding 500ms of spiking activity for all M1 units, binned at 100ms, were used as predictors. A model was also fit on baseline, non-reaching, neural trajectories, calculated by projecting DLS spiking activity from five seconds to four second before pellet touch onto each of the first two computed principal components. The predictive ability of these models was assessed by calculating the correlation between the actual neural trajectories and the predicted trajectories.

### NREM rhythm detection (Figure 4d)

The NREM rhythm detection applied here is based on an algorithm we have developed previously (Kim et al., 2019, Silversmith, et al., 2020). A mean LFP channel was generated in M1 for NREM rhythm classification by averaging across all channels (same as used for sleep classification). To detect sleep spindles, this mean signal was filtered in the spindle band (10 – 16 Hz) using a zero-phase shifted, third order Butterworth filter. A smoothed envelope was calculated by computing the magnitude of the Hilbert transform of this signal then convolving it with a Gaussian window. Next, we determined two upper thresholds for spindle detection based on the mean and standard deviation (s.d.) of the spindle band envelope during NREM. Epochs in which the spindle envelope exceeded 2.5 s.d. above the mean for at least one sample and the spindle power exceeded 1.5 s.d. above the mean for at least 500ms were detected as spindles. Then, spindles that were sufficiently close in time (<300 ms) were combined. To detect slow oscillations and delta waves, the mean M1 signal was filtered in a low frequency band (2^nd^ order, zero phase shifted, high pass Butterworth filter with a cutoff at 0.1Hz followed by a 5^th^ order, zero phase shifted, low pass Butterworth filter with a cutoff at 4Hz). Next, all positive-to-negative zero crossings during NREM were identified, along with the previous peaks, the following troughs, and the surrounding negative-to-positive zero crossings. Each identified epoch was considered a slow oscillation if the peak was in the top 15% of peaks, the trough was in the top 40% of troughs and the time between the negative-to-positive zero crossings was greater than 300ms but did not exceed 1 second. Each identified epoch was considered a delta wave if the peak was in the bottom 85% of peaks, the trough was in the top 40% of troughs and the time between the negative-to-positive zero crossings was greater than 250ms.

### Characterizing putatively monosynaptically connected M1 and DLS units (Figure 4b; Supplemental Figure 2b)

We characterized putatively monosynaptically connected pairs of M1 and DLS units by calculating the cross correlation of spiking activity binned at 1ms during the first five minutes of NREM during pre- and post-sleep concatenated together (10 minutes total) on each day of training for each pair of M1 and DLS units. We then measured the mean value of the short-latency cross correlation for each pair (1-10ms time lag centered on DLS spiking; consistent with the conduction and synaptic delay between M1 and DLS; Koralek et al., 2013) and compared this value to a shuffled distribution generated by shuffling DLS spike time bins and recalculating the cross correlation 1,000 times. If the non-shuffled short-latency correlation magnitude was greater than 95% of the shuffled distribution values, we classified the pair of units as putatively connected.

### Comparing corticostriatal transmission strength across behavioral states (Figure 4c; Supplemental Figure 3a&b)

To compare corticostriatal transmission strength across behavioral states, we generated a cross correlation of spiking activity binned at 1ms from each behavioral state (NREM, REM, and wake) for all putatively connected pairs of M1 and DLS units, during both pre- and post-sleep. To account for firing rate differences across states, each pair’s cross correlation was normalized by subtracting the mean cross correlation values from 100-150ms time lag.

### Comparing corticostriatal transmission strength across NREM rhythms (Figure 4g; Supplemental Figure 3c&d)

To compare corticostriatal transmission strength across NREM rhythms, we generated a cross correlation of spiking activity binned at 1ms from each NREM rhythm (sleep spindles, delta waves, and slow oscillations) for all putatively connected pairs of M1 and DLS units. Spiking during sleep spindles consisted of spiking during the one second centered on sleep spindle peak (−500ms to 500ms). Spiking during slow oscillations and delta waves consisted of spiking during the one second around upstate peak (−500ms to 500ms). To account for the influence of firing rate differences or changes in LFP-phase locking across NREM rhythms, we applied a normalization step we previously developed (Silversmith, et al., 2020). Briefly, we generated shuffled cross correlations between each M1 and DLS unit pair, with DLS spike times shuffled with respect to the NREM rhythm in which it fired. In this approach, both units maintain all their first-order relationships with the NREM rhythm; for example, the number of spikes, phase locking values, and phase preferences of individual units do not change after shuffling. However, the shuffling breaks the statistical relationship between the two neurons under examination. We repeated this shuffling 25 times and then subtracted the mean shuffled cross correlation from the unshuffled cross correlation.

### NREM rhythm modulation (Figure 4e&f; Supplemental Figure 4)

To determine the sleep spindle modulation of individual M1 and DLS units, spiking during each sleep spindle was time locked to the peak of the filtered LFP and binned at 10ms. Spiking was averaged across sleep spindles and modulation was calculated by taking the minimum to maximal firing rate bin in the second around sleep spindle peak (−500ms to 500ms) divided by the minimum to maximal firing rate bin in a second long baseline period before each spindle (−1500ms to −500ms relative to spindle peak). To determine slow oscillation and delta wave modulation of individual M1 and DLS units, spiking during each slow oscillation or delta wave was time locked to the peak of the upstate and binned at 10ms. Spiking was averaged across slow oscillations or delta waves and modulation was calculated by taking the minimum to maximal firing rate bin in the second around upstate peak (−500ms to 500ms) divided by the minimum to maximal firing rate bin in a second long baseline period before each slow oscillation or delta wave (−1500ms to −500ms relative to upstate peak).

### Characterizing reach modulated (RM) units (Figure 5; Supplemental Figure 5; Supplemental Figure 6)

To characterize M1 and DLS reach modulated units, we generated trial-averaged peri-event time histograms (PETHs) of spiking activity for individual units during reaching locked to pellet touch in 25ms bins, from 5 seconds before to 5 seconds after pellet touch (400 total bins). Each unit’s PETH was then z-scored and reach modulation was measured by taking the sum of the absolute value of the time bins from 1 second before pellet touch to 1 second after pellet touch (80 total bins). We then generated a distribution of shuffled modulations by shuffling all time bins and recalculating the modulation of the shuffled PETH and repeating this shuffling procedure one thousand times. Units with a non-shuffled modulation greater than the 99% percentile of the shuffled distribution were considered significantly reach modulated.

### Characterizing DLS units with strong or weak M1 reach modulated input (Figure 5e&f; Supplemental Figure 5e&f; Supplemental Figure 6e&f)

To characterize DLS units with strong or weak M1 reach modulated input, we calculated the number of reach modulated M1 units that were putatively connected to each DLS unit. If a DLS unit was connected to 3 or more reach modulated M1 units, we classified that DLS unit as having strong M1 reach modulated input, if a DLS unit was connected to 2 or less M1 reach modulated units, we classified that DLS unit as having weak or no M1 reach modulated input.

### Measuring corticostriatal transmission strength changes within pre- and post-sleep (Figure 6; Supplemental Figure 8)

To measure changes in corticostriatal transmission strength within pre- and post-sleep, we generated a cross correlation of spiking activity binned at 1ms from NREM activity during the first and second half of pre- and post-sleep. This was done for two populations of M1 and DLS unit pairs. The first population was all M1 and DLS unit pairs that contained a RM M1 unit, a DLS unit that was putatively connected to a RM M1 unit, and contained both M1 and DLS units that were significantly modulated to sleep spindles. The second population was all M1 and DLS unit pairs that contained a RM M1 unit, a DLS unit that was putatively connected to a RM M1 unit, and contained M1 and DLS units that were both not significantly modulated to sleep spindles. To determine which units were modulated to sleep spindles, we generated perievent time histograms (PETHs) of sleep spindle activity locked to spindle peak in 10ms bins from 2 seconds before to 2 second after spindle peak (400 bins), averaged across all spindles. Sleep spindle modulation was then calculated by taking the minimum to maximal firing rate bin within the 1 second period centered on spindle peak (−500ms to 500ms). We then generated a distribution of shuffled modulations by shuffling the time bins and recalculating the modulation of this shuffled PETH. This shuffling procedure was repeated one thousand times to generate a distribution. Units with a non-shuffled modulation greater than the 99% percentile of the shuffled distribution were considered significantly sleep spindle modulated.

### Sleep spindle and slow oscillation proximity (Figure 7)

Slow oscillation to sleep spindle proximity was determined by measuring the temporal proximity of the preceding slow oscillation zero-crossing (positive to negative LFP) to each sleep spindle peak. To determine the influence of slow oscillation proximity on sleep spindle modulation, we generated two PETHs locked to spindle peak and binned at 10ms for each unit. The first PETH was generated with sleep spindles that had a preceding slow oscillation within 500ms (“nested spindles”) and the second PETH was generated with sleep spindles that did not have a preceding slow oscillation within 500ms (“isolated spindles”). As there were more isolated spindles than nested spindles, the number of events used to generate each PETH was matched by randomly selecting isolated spindles to match the number of nested spindles. Modulation was then assessed by determining the peak firing rate bin in each PETH.

## References

1. Isoda, M. & Hikosaka, O. Cortico-basal ganglia mechanisms for overcoming innate, habitual and motivational behaviors. Eur. J. Neurosci. 33, 2058–2069 (2011).

2. Smith, K. S. & Graybiel, A. M. Habit formation. Dialogues Clin. Neurosci. 18, 33–43 (2016).

3. Robbins, T. W. & Costa, R. M. Habits. Current Biology (2017). doi:10.1016/j.cub.2017.09.060

4. Graybiel, A. M. Habits, Rituals, and the Evaluative Brain. Annu. Rev. Neurosci. 31, 359–387 (2008).

5. Yin, H. H. & Knowlton, B. J. The role of the basal ganglia in habit formation. Nat. Rev. Neurosci. 7, 464–476 (2006).

6. Costa, R. M., Cohen, D. & Nicolelis, M. A. L. Differential Corticostriatal Plasticity during Fast and Slow Motor Skill Learning in Mice. Curr. Biol. 14, 1124–1134 (2004).

7. Neely, R. M., Koralek, A. C., Athalye, V. R., Costa, R. M. & Carmena, J. M. Volitional Modulation of Primary Visual Cortex Activity Requires the Basal Ganglia. Neuron 0, (2018).

8. Lemke, S. M., Ramanathan, D. S., Guo, L., Won, S. J. & Ganguly, K. Emergent modular neural control drives coordinated motor actions. Nat. Neurosci. 22, 1122–1131 (2019).

9. Koralek, A. C., Costa, R. M. & Carmena, J. M. Temporally Precise Cell-Specific Coherence Develops in Corticostriatal Networks during Learning. Neuron 79, 865–872 (2013).

10. Santos, F. J., Oliveira, R. F., Jin, X. & Costa, R. M. Corticostriatal dynamics encode the refinement of specific behavioral variability during skill learning. Elife 4, (2015).

11. Dang, M. T. et al. Disrupted motor learning and long-term synaptic plasticity in mice lacking NMDAR1 in the striatum. Proc. Natl. Acad. Sci. U. S. A. (2006). doi:10.1073/pnas.0601758103

12. Koralek, A. C., Jin, X., Long II, J. D., Costa, R. M. & Carmena, J. M. Corticostriatal plasticity is necessary for learning intentional neuroprosthetic skills. Nature 483, 331–335 (2012).

13. Calabresi, P., Pisani, A., Mercuri, N. B. & Bernardi, G. Long-term Potentiation in the Striatum is Unmasked by Removing the Voltage-dependent Magnesium Block of NMDA Receptor Channels. Eur. J. Neurosci. (1992). doi:10.1111/j.1460-9568.1992.tb00119.x

14. Charpier, S. & Deniau, J. M. In vivo activity-dependent plasticity at cortico-striatal connections: Evidence for physiological long-term potentiation. Proc. Natl. Acad. Sci. U. 5. A. (1997). doi:10.1073/pnas.94.13.7036

15. Yang, G. et al. Sleep promotes branch-specific formation of dendritic spines after learning. Science (80-.). 344, 1173–1178 (2014).

16. Ramanathan, D. S., Gulati, T. & Ganguly, K. Sleep-Dependent Reactivation of Ensembles in Motor Cortex Promotes Skill Consolidation. PLOS Biol. 13, e1002263 (2015).

17. Gulati, T., Ramanathan, D. S., Wong, C. C. & Ganguly, K. Reactivation of emergent task-related ensembles during slow-wave sleep after neuroprosthetic learning. Nat. Neurosci. 17, 1107–1113 (2014).

18. Kim, J., Gulati, T. & Ganguly, K. Competing Roles of Slow Oscillations and Delta Waves in Memory Consolidation versus Forgetting. Cell 179, 514–526.e13 (2019).

19. Vahdat, S., Fogel, S., Benali, H. & Doyon, J. Network-wide reorganization of procedural memory during NREM sleep revealed by fMRI. Elife 6, (2017).

20. Boutin, A. et al. Transient synchronization of hippocampo-striato-thalamo-cortical networks during sleep spindle oscillations induces motor memory consolidation. Neuroimage 169, 419–430 (2018).

21. Doyon, J., Gabitov, E., Vahdat, S., Lungu, O. & Boutin, A. Current issues related to motor sequence learning in humans. Curr. Opin. Behav. Sci. 20, 89–97 (2018).

22. Doyon, J. & Benali, H. Reorganization and plasticity in the adult brain during learning of motor skills. Curr. Opin. Neurobiol. 15, 161–167 (2005).

23. Klinzing, J. G., Niethard, N. & Born, J. Mechanisms of systems memory consolidation during sleep. Nat. Neurosci. 22, 1598–1610 (2019).

24. Rasch, B. & Born, J. About Sleep’s Role in Memory. Physiol. Rev. 93, 681–766 (2013).

25. Squire, L. R., Genzel, L., Wixted, J. T. & Morris, R. G. Memory consolidation. Cold Spring Harb. Perspect. Biol. 7, a021766 (2015).

26. Lipton, D. M., Gonzales, B. J. & Citri, A. Dorsal Striatal Circuits for Habits, Compulsions and Addictions. Front. Syst. Neurosci. 13, 28 (2019).

27. Lee, J. L. C., Nader, K. & Schiller, D. An Update on Memory Reconsolidation Updating. Trends Cogn. Sci. 21, 531–545 (2017).

28. Egert, D. G. et al. Cellular-scale silicon probes for high-density, precisely-localized neurophysiology. J. Neurophysiol. (2020). doi:10.1152/jn.00352.2020

29. Aoki, S. et al. An open cortico-basal ganglia loop allows limbic control over motor output via the nigrothalamic pathway. Elife 8, (2019).

30. Watson, B. O., Levenstein, D., Greene, J. P., Gelinas, J. N. & Buzsáki, G. Network Homeostasis and State Dynamics of Neocortical Sleep. Neuron 90, 839–852 (2016).

31. Rueda-Orozco, P. E. & Robbe, D. The striatum multiplexes contextual and kinematic information to constrain motor habits execution. Nat. Neurosci. 18, 453–460 (2015).

32. Flint, R. D., Scheid, M. R., Wright, Z. A., Solla, S. A. & Slutzky, M. W. Long-Term Stability of Motor Cortical Activity: Implications for Brain Machine Interfaces and Optimal Feedback Control. J. Neurosci. 36, 3623 (2016).

33. Yazdan-Shahmorad, A., Silversmith, D. B., Kharazia, V. & Sabes, P. N. Targeted cortical reorganization using optogenetics in non-human primates. Elife 7, (2018).

34. Thorn, C. A. & Graybiel, A. M. Differential Entrainment and Learning-Related Dynamics of Spike and Local Field Potential Activity in the Sensorimotor and Associative Striatum. J. Neurosci. 34, 2845–2859 (2014).

35. Huber, R., Felice Ghilardi, M., Massimini, M. & Tononi, G. Local sleep and learning. Nature 430, 78–81 (2004).

36. Durkin, J. et al. Cortically coordinated NREM thalamocortical oscillations play an essential, instructive role in visual system plasticity. Proc. Natl. Acad. Sci. 114, 10485–10490 (2017).

37. Ramanathan, D. S., Gulati, T. & Ganguly, K. Sleep-Dependent Reactivation of Ensembles in Motor Cortex Promotes Skill Consolidation. PLOS Biol. 13, e1002263 (2015).

38. Silversmith, D. B., Lemke, S. M., Egert, D., Berke, J. D. & Ganguly, K. The Degree of Nesting between Spindles and Slow Oscillations Modulates Neural Synchrony. J. Neurosci. 40, 4673–4684 (2020).

39. Niethard, N., Ngo, H.-V. V., Ehrlich, I. & Born, J. Cortical circuit activity underlying sleep slow oscillations and spindles. Proc. Natl. Acad. Sci. 115, E9220–E9229 (2018).

40. Hoffman, K. L. & McNaughton, B. L. Coordinated reactivation of distributed memory traces in primate neocortex. Science (80-.). 297, 2070–2073 (2002).

41. Lansink, C. S., Goltstein, P. M., Lankelma, J. V., McNaughton, B. L. & Pennartz, C. M. A. Hippocampus Leads Ventral Striatum in Replay of Place-Reward Information. PLoS Biol. 7, e1000173 (2009).

42. Sjulson, L., Peyrache, A., Cumpelik, A., Cassataro, D. & Buzsáki, G. Cocaine Place Conditioning Strengthens Location-Specific Hippocampal Coupling to the Nucleus Accumbens. Neuron 98, 926–934.e5 (2018).

43. Ji, D. & Wilson, M. A. Coordinated memory replay in the visual cortex and hippocampus during sleep. Nat. Neurosci. 10, 100–107 (2007).

44. Girardeau, G., Inema, I. & Buzsáki, G. Reactivations of emotional memory in the hippocampus-amygdala system during sleep. Nat. Neurosci. 20, 1634–1642 (2017).

45. Ribeiro, S. et al. Long-Lasting Novelty-Induced Neuronal Reverberation during Slow-Wave Sleep in Multiple Forebrain Areas. PLoS Biol. 2, e24 (2004).

46. Rasch, B. & Born, J. About Sleep’s Role in Memory. Physiol. Rev. 93, 681–766 (2013).

47. Yu, J. Y., Liu, D. F., Loback, A., Grossrubatscher, I. & Frank, L. M. Specific hippocampal representations are linked to generalized cortical representations in memory. Nat. Commun. (2018). doi:10.1038/s41467-018-04498-w

48. Tingley, D. & Peyrache, A. On the methods for reactivation and replay analysis. Philosophical Transactions of the Royal Society B: Biological Sciences (2020). doi:10.1098/rstb.2019.0231

49. Yin, H. H. et al. Dynamic reorganization of striatal circuits during the acquisition and consolidation of a skill. 12, (2009).

50. Rosanova, M. & Ulrich, D. Pattern-specific associative long-term potentiation induced by a sleep spindle-related spike train. J. Neurosci. 25, 9398–9405 (2005).

51. Chauvette, S., Seigneur, J. & Timofeev, I. Sleep oscillations in the thalamocortical system induce long-term neuronal plasticity. Neuron 75, 1105 (2012).

52. Churchland, M. M. et al. Neural population dynamics during reaching. Nature 487, 51–6 (2012).

53. Fischer, S., Hallschmid, M., Elsner, A. L. & Born, J. Sleep forms memory for finger skills. Proc. Natl. Acad. Sci. 99, 11987–11991 (2002).

54. Walker, M. P., Brakefield, T., Morgan, A., Hobson, J. A. & Stickgold, R. Practice with Sleep Makes Perfect: Sleep-Dependent Motor Skill Learning. Neuron 35, 205–211 (2002).

55. Nagai, H. et al. Sleep consolidates motor learning of complex movement sequences in mice. Sleep (2017). doi:10.1093/sleep/zsw059

56. Gerdeman, G. L., Partridge, J. G., Lupica, C. R. & Lovinger, D. M. It could be habit forming: drugs of abuse and striatal synaptic plasticity. Trends Neurosci. 26, 184–192 (2003).

## Methods References

1. Wong, C. C., Ramanathan, D. S., Gulati, T., Won, S. J. & Ganguly, K. An automated behavioral box to assess forelimb function in rats. J. Neurosci. Methods 246, 30–7 (2015).

2. Mathis, A. et al. DeepLabCut: markerless pose estimation of user-defined body parts with deep learning. Nat. Neurosci. 21, 1281–1289 (2018).

3. Watson, B. O., Levenstein, D., Greene, J. P., Gelinas, J. N. & Buzsáki, G. Network Homeostasis and State Dynamics of Neocortical Sleep. Neuron 90, 839–852 (2016).

4. J, K., T, G. & K, G. Competing Roles of Slow Oscillations and Delta Waves in Memory Consolidation versus Forgetting. Cell 179, (2019).

5. Silversmith, D. B., Lemke, S. M., Egert, D., Berke, J. D. & Ganguly, K. The Degree of Nesting between Spindles and Slow Oscillations Modulates Neural Synchrony. J. Neurosci. 40, 4673–4684 (2020).

6. Koralek, A. C., Costa, R. M. & Carmena, J. M. Temporally Precise Cell-Specific Coherence Develops in Corticostriatal Networks during Learning. Neuron 79, 865–872 (2013).

